# Age-elevated prostaglandin E_2_ enhances mortality to influenza infection

**DOI:** 10.1101/2021.12.02.470775

**Authors:** Judy Chen, Jane C. Deng, Rachel Zemans, Min Zhang, Marc Peters-Golden, Daniel R. Goldstein

## Abstract

Aging impairs the immune responses to influenza A virus (IAV), resulting in increased mortality to IAV infections in older adults. With aging, there is reduced number and impaired function of alveolar macrophages (AMs), cells critical for defense against IAV. However, factors within the aged lung that impair AMs are not fully known. Using a murine model of IAV infection, we observed that aging increased the level of prostaglandin E_2_ (PGE_2_) in the bronchoalveolar lavage fluid (BALF) of aged mice compared to young mice. Blockade of the PGE_2_ receptor EP2 in aged mice increased AM numbers and subsequently enhanced survival to IAV. Additionally, PGE_2_ impaired the mitochondrial health of AMs. We also identified senescent type II alveolar epithelial cells (AECs) as a source of the aged-associated PGE_2_ in the lung. Our results reveal a crosstalk between AECs and AMs, via PGE_2_, that compromises host defense to IAV infection with aging.

## Introduction

Influenza A virus (IAV) remains a serious health burden that disproportionally affects older individuals. While most people who contract IAV only experience mild to moderate symptoms, older individuals (≥65 years of age) experience higher rates of susceptibility, mortality, and complications such as secondary bacterial infections (Iuliano et al., 2018; Thompson et al., 2004). For example, 86% of the deaths due to 2017-2018 seasonal IAV infections in the United States occurred in individuals 65 years or older (Centers for Disease Control and Prevention, 2019), even though this age group is only 13% of the total US population (United States Census Bureau, 2011). This susceptibility is of increasing concern as the average age of the world’s population continues to increase, especially in developed countries (United Nations, 2020).

Currently, there are three classes of anti-IAV medications: adamantanes, neuraminidase inhibitors, and cap-dependent endonuclease inhibitors (Cooper et al., 2003; Uyeki et al., 2019). However, these drugs show limited effectiveness in older individuals and the emergence of IAV strains that are resistant to these treatments will limit their future use (Cooper et al., 2003; Uyeki et al., 2019). Given the limitations of the current therapies and the increasing burden of IAV in older adults, we need to understand the consequences of aging on the immune response against IAV so that we can develop novel age-specific therapies.

We have previously examined the effects of aging on alveolar macrophages (AMs), the sentinel airspace and airway resident macrophages that are the first responders to respiratory pathogens and play a key role in maintaining homeostasis of the lungs. The importance of AMs in the control of an IAV infection is best evidenced by the rapid weight loss, increased tissue damage, and poor survival in murine models of genetic AM deficiency and pharmacological AM depletion (Cardani et al., 2017; Wong et al., 2017). Aging impairs AM functions such as phagocytosis, efferocytosis, scavenger receptor expression and proliferation, and is associated with significant transcriptional changes (McQuattie-Pimentel et al., 2021; Wong et al., 2017). Adoptive transfer experiments in which AMs isolated from young mice are transferred into aged mice and vice versa show that the age-associated transcriptomic differences in AMs are largely driven by the local aged lung microenvironment rather than cell-intrinsic factors (McQuattie-Pimentel et al., 2021). However, the factors within the aged lung microenvironment that contribute to the age-associated defects of AMs remain largely undefined.

The lipid prostaglandin E_2_ (PGE_2_) is a prostanoid with pleotropic functions in the regulation of immune cells but with unclear implications for respiratory health and host defense with aging. PGE_2_ is produced from the synthesis of prostaglandin H_2_ (PGH_2_) from arachidonic acid by the cyclooxygenase (COX)-1 and COX-2 proteins and the further conversion of PGH_2_ by the PGE_2_ synthases: microsomal PGE_2_ synthase-1 (mPGES-1), mPGES-2, and cytosolic PGE_2_ synthase (cPGES) (Nakanishi and Rosenberg, 2013; Ricciotti and FitzGerald, 2011). PGE_2_ signals through four G-protein coupled receptors: EP1, EP2, EP3, and EP4 (Nakanishi and Rosenberg, 2013; Ricciotti and FitzGerald, 2011). Of these receptors, EP2 and EP4 have been shown to be upregulated on AMs during IAV infection (Coulombe et al., 2014). Furthermore, PGE_2_ signaling through the EP2 receptor regulates AM functions such as phagocytosis of bacteria (Aronoff et al., 2004), toll-like receptor expression (Degraaf et al., 2014), and production of suppressor of cytokine signaling 3 (SOCS3) (Speth et al., 2016). Prior studies, including our own, have found that PGE_2_ is increased in the lung with aging in mice before infection (Penke et al., 2020; Vijay et al., 2015). In addition to aging, PGE_2_ levels are known to increase with various inflammatory conditions such as rheumatoid arthritis (McCoy et al., 2002) and cancer (Nakanishi and Rosenberg, 2013), as well as infections such as IAV infection (Coulombe et al., 2014). But whether PGE_2_ is a dominant microenvironmental factor in the aged lung that compromises the AM response to viral function is not known.

Cellular senescence, commonly referred to as senescence, is a hallmark of aging (López-Otín et al., 2013). Senescence is thought to be due to the accumulation of DNA damage, telomere shortening, and/or mitochondrial dysfunction (Di Micco et al., 2021; Kumari and Jat, 2021; López-Otín et al., 2013). One aspect of senescence is the senescence-associated secretory phenotype (SASP), by which senescent cells secrete inflammatory mediators such as cytokines, chemokines, and growth factors, at steady state (Di Micco et al., 2021; Kumari and Jat, 2021). As senescent cells accumulate with age, increased production of SASP factors contribute to inflammaging, the low-grade, chronic, sterile inflammation associated with aging (Franceschi and Campisi, 2014; Franceschi et al., 2018).

Here, we employed murine models and show that PGE_2_ levels increased within the lung alveolar space with aging. We revealed that PGE_2_,through the EP2 receptor, limits AM proliferation and mitochondrial fitness with aging, and most importantly PGE_2_ reduces survival of aged mice to lethal IAV infection. Additionally, we identified senescent type II alveolar epithelial cells (AECs) to be a contributor to age-associated elevations of PGE_2_ in the lung. Our study has revealed a pathophysiological relationship between type II AECs and AMs via elevated PGE_2_ that impairs host defense to IAV with aging.

## Results

### Aging increases PGE_2_ levels in the lung before and after IAV infection

To begin to interrogate the role of PGE_2_ in respiratory health and host defense with aging, we first measured PGE_2_ levels in the plasma of young (2-4 months) and aged (18-22 months) female non-infected C57BL/6 mice by ELISA. We observed a ∼2-fold increase in PGE_2_ concentration in aged mice as compared to young mice (Supplemental Figure 1A), confirming that aging indicates that aging leads to increased systemic levels of PGE_2_.

Next, we measured levels of PGE_2_ within the bronchoalveolar lavage fluid (BALF) of young and aged female non-infected C57BL/6 mice. We observed a ∼2-fold increase of PGE_2_ in the BALF of aged mice compared to young mice (Figure 1A). To confirm that this finding is not genotype- or sex-specific, we also measured PGE_2_ in the BALF of young (6 months) and aged (22 months) male non-infected UM-HET3 mice, a 4-way crossed outbred mouse strain used by the National Institute on Aging Interventions Testing Program (Miller et al., 2007). Similar to the female C57BL/6 mice, the aged male UM-HET3 exhibited an ∼3-fold increase of PGE_2_ within the BALF compared to the young mice (Figure 1B).

**Figure 1:**
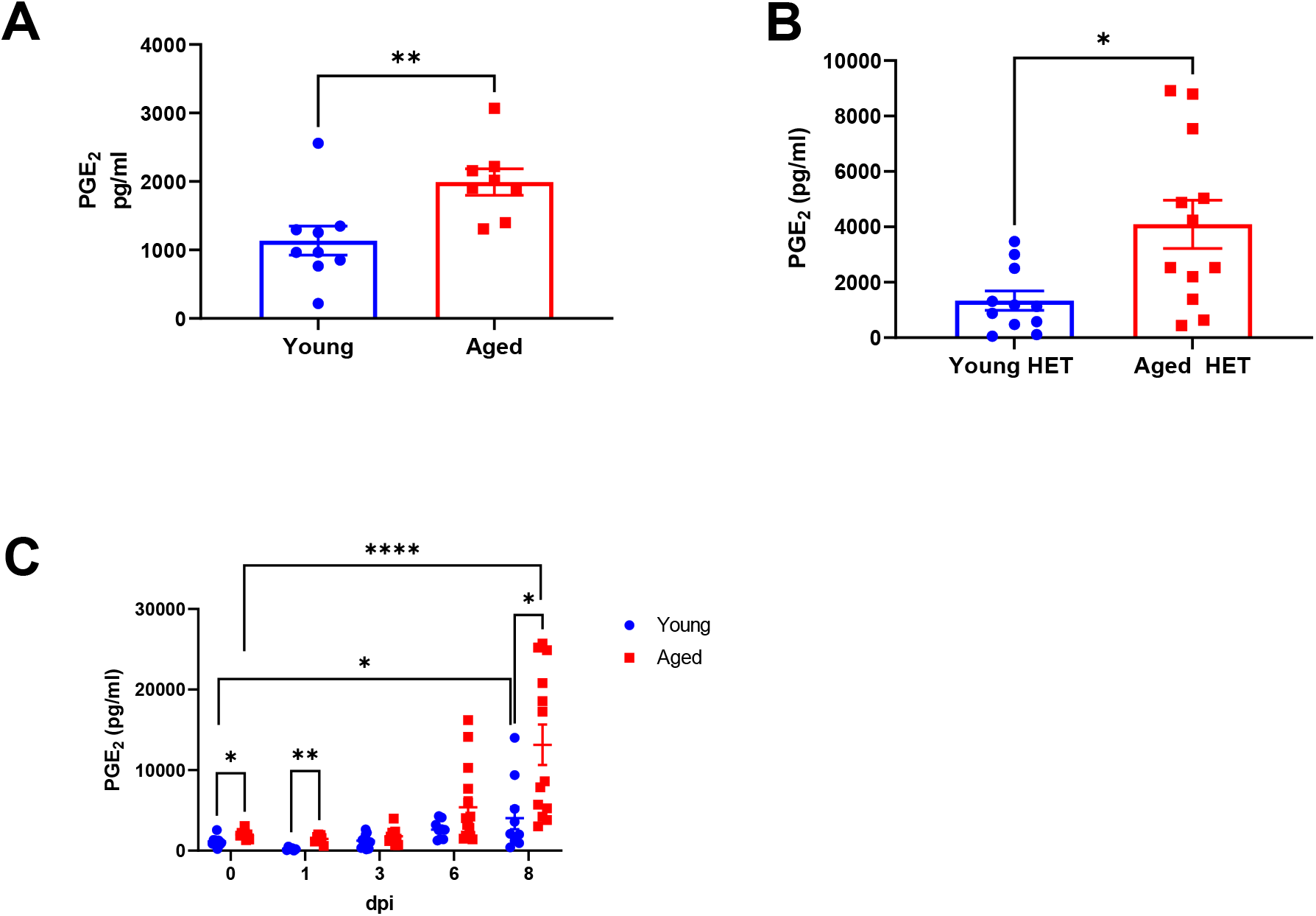
Aging increases PGE_2_ levels in the lung before and after IAV infection. (**A-B**) PGE_2_ measured from the BALF of non-infected and infected young and aged mice. (**A**) PGE_2_ levels in the BALF of young (2-4 months) and aged (18-22 months) non-infected female C57BL/6 mice. Data analyzed by Mann-Whitney. (**B**) PGE_2_ levels in the BALF of young (6 months) and aged (22 months) non-infected male UM-HET3 mice. Data analyzed by Mann-Whitney. (**C**) PGE_2_ measured from the BALF of young (2-4 months) and aged (18-22 months) female C57BL/6 mice infected with 400 pfu of PR8 IAV intranasally at 0, 1, 3, 6 and 8 days post infection (dpi). The PGE_2_ measurements at 0 DPI is duplicated data from Fig 1A. Data analyzed by Mann-Whitney with the Benjamini, Krieger and Yekutieli method to correct for multiple comparisons. Error bars represent SEM. Each point represents a biological replicate. * p< 0.05, ** p< 0.005, ****p<0.0001.

How aging impacts PGE_2_ levels within the BALF during respiratory viral infections, in particular IAV, remains to be elucidated. To examine this, we infected young and aged female C57BL/6 mice with 400 plaque forming units (pfu) of IAV intranasally (i.n.) as 400 pfu is the lethal dose 70% (LD_70_) in aged mice (Supplemental Figure 1B, survival tracked up to 15 days post infection (dpi)). Note, for all subsequent infections, female mice were used and infected with 400 pfu, unless otherwise stated). The BALF was then collected from the mice at 0, 1, 3, 6, and 8 dpi. IAV infection increased PGE_2_ levels within the BALF in both young and aged mice (Figure 1C). Notably, aged mice exhibited ∼4-5 -fold higher levels of PGE_2_ within the BALF as compared to young mice by 8 dpi. Overall, these results indicate that PGE_2_ levels increase after IAV infection, and this increase is exacerbated by aging.

### Blocking PGE_2_ signaling via the EP2 receptor increases AM numbers in aged mice

Aging reduces the number of AMs in female C57BL/6 mice (Wong et al., 2017). We confirmed this age-associated phenotype is not sex- or strain-specific by enumerating AMs in the BALF of young and aged male UM-HET3 mice (Supplemental Figure 2A).

We recently showed that PGE_2_ limits AM proliferation *in vitro* through the PGE_2_ receptor EP2 (Penke et al., 2020). To investigate whether PGE_2_ signaling affects AM numbers *in vivo,* we blocked PGE_2_ signaling *in vivo* via intraperitoneal (i.p.) injections of an antagonist against the EP2 receptor (af Forselles et al., 2011). This treatment reduced PGE_2_-induced cAMP in the lungs (Supplemental Figure 2B), validating our approach. Non-infected aged C57BL/6 mice were i.p. injected daily with the EP2 antagonist or vehicle control for 7 days. We collected the BALF and enumerated AMs (defined as CD45^+^ CD11c^+^ SiglecF^+^) by flow cytometry. Blocking PGE_2_ signaling through the EP2 receptor increased total AM numbers by ∼1.3-fold in aged mice (Figure 2A). In contrast, blocking the EP2 receptor in young C57BL/6 mice failed to alter total AM numbers (Supplemental Figure 2C). These results suggest that PGE_2_ signaling becomes aberrant with aging and limits AM numbers.

**Figure 2:**
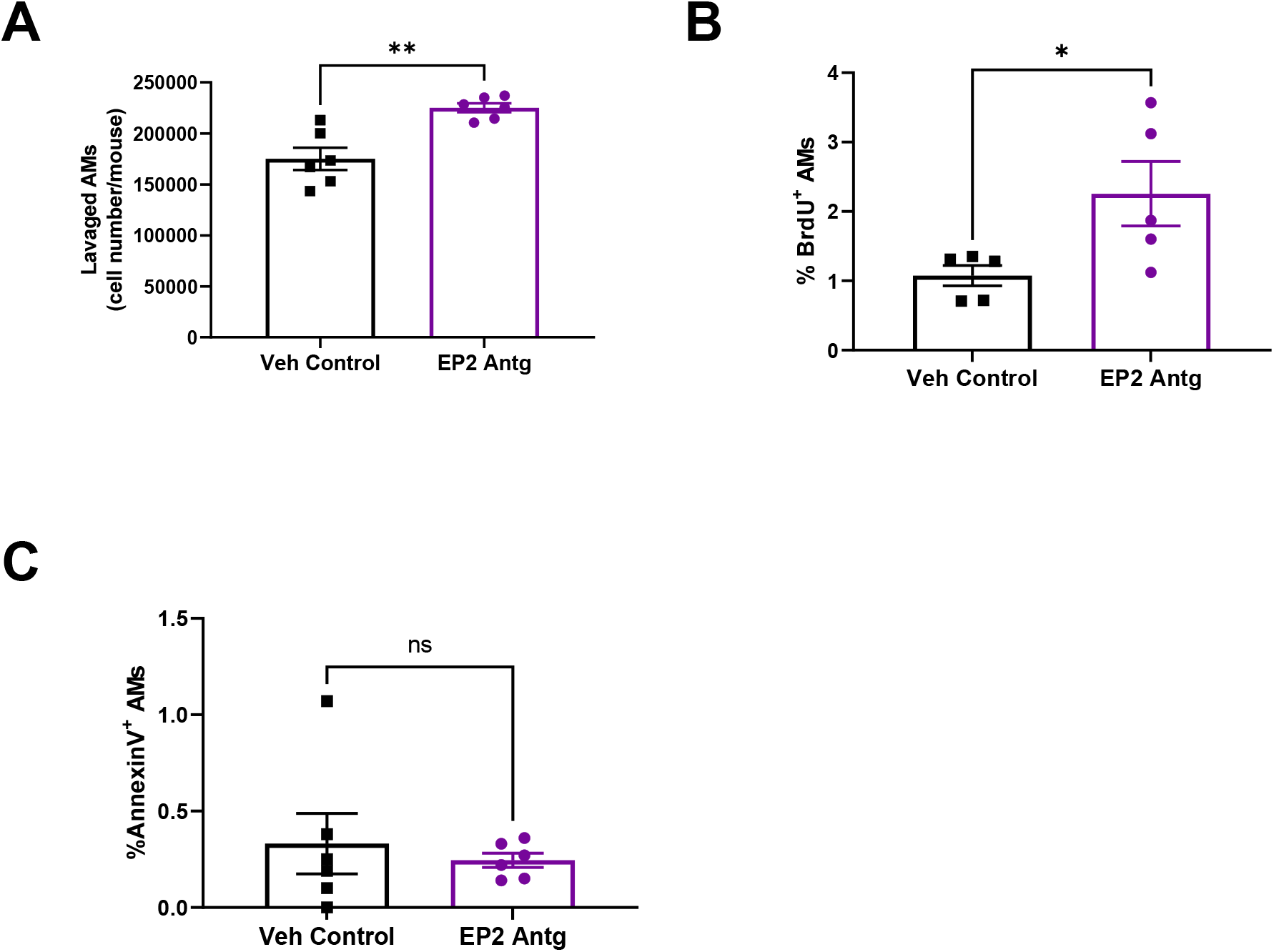
Blocking PGE_2_ signaling via the EP2 receptor increases AM numbers in aged mice. Aged (18-22 months) C57BL/6 mice were given 7 daily i.p. injections of 10mg/kg EP2 antagonist. Concurrently, BrdU (0.8 mg/ml) was given in the drinking water *ad libitum.* AMs (i.e., CD45^+^ CD11c^+^ SiglecF^+^) were then collected through BAL and analyzed by flow cytometry for BrdU incorporation and Annexin V staining. (**A**) Total cell count of lavaged AMs. (**B**) Percentage of AMs that are BrdU^+^. (**C**) Percentage of AMs that are Annexin V^+^. Statistical significance analyzed by Mann-Whitney. Error bars represent SEM. Each point represents a biological replicate. * p< 0.05, ** p< 0.005.

Increased proliferation and/or reduced apoptosis could explain the increase in AM number. To determine if blocking the PGE_2_ receptor EP2 increases AM cell numbers through proliferation, aged mice treated with the EP2 antagonist or vehicle control were also administered BrdU, a thymidine analog that is incorporated into newly synthesized DNA of proliferating cells (Cameron, 2006). Flow cytometry analysis revealed that blocking the EP2 receptor in aged mice led to a ∼2-fold increase in the percent of BrdU positive AMs compared to the control (Figure 2B). To assess if blocking PGE_2_ signaling alters apoptosis of AMs, we stained AMs from aged mice treated with EP2 antagonist or vehicle control with Annexin V and analyzed via flow cytometry. Our results indicated that EP2 receptor blockade does not affect AM apoptosis (Figure 2C). Overall, these results show that PGE_2_ limits AM numbers with aging, via the EP2 receptor, by reducing AM proliferation and without altering apoptosis.

To determine if aging alters the expression of the EP2 receptor on AMs, we re-analyzed a publicly available RNA-seq dataset (GEO GSE134397) of sorted AMs from young and aged mice (McQuattie-Pimentel et al., 2021). The trimmed mean of M-values (TMM) normalized counts from the RNA-seq dataset showed no age-associated differences in expression of the EP2 receptors on AMs on a transcriptomic level (Supplemental Figure 2D). This indicates that the differential response of EP2 antagonism in young versus aged mice is independent of altered expression of the EP2 receptor.

### Prophylactic blockade of PGE_2_ signaling through the EP2 receptor improves survival to IAV in aged mice

AMs are crucial for host defense against respiratory viruses such as IAV (Cardani et al., 2017; Wong et al., 2017). Our results suggest that PGE_2_ limits AM numbers in non-infected aged mice. Therefore, we hypothesized that blocking PGE_2_ signaling in aged mice prior to infection and boosting AM numbers would subsequently increase survival to lethal IAV infection. Hence, we gave aged mice the EP2 receptor antagonist prophylactically for 7 days to increase AM numbers (as was demonstrated in Figure 2A), followed by an i.n., infection with IAV (Figure 3A). We found that EP2 antagonist treatment significantly increased survival in aged IAV infected mice from ∼20% (vehicle control) to ∼50% (EP2 antagonist) (Figure 3B, survival tracked up to 18 dpi).

**Figure 3:**
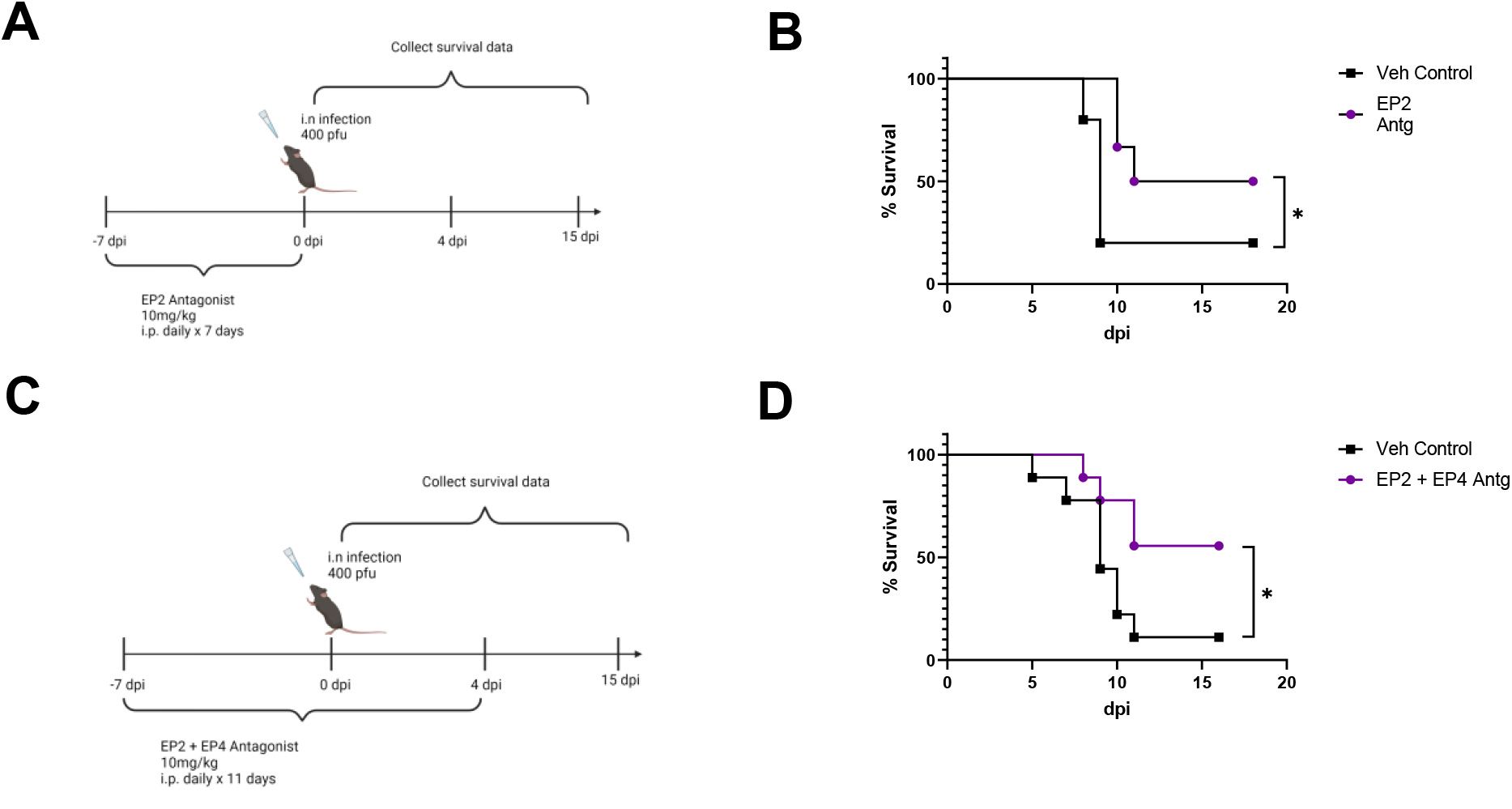
Prophylactic blockade of PGE_2_ signaling through the EP2 receptor improves survival to IAV in aged mice. Aged (18-22 months) female C57BL/6 mice were given 7 daily i.p. injections of 10mg/kg of the EP2 antagonist starting 7 days before infection. The mice were then infected with 400 pfu of PR8 IAV intranasally as depicted in (**A**). Survival of mice (**B**) was tracked daily. n= 5-6 / group. Aged (18-22 months) female C57BL/6 mice were given 11 daily i.p. injections of 10mg/kg of the EP2 and EP4 antagonist from days -7 pre-infection to 4 dpi. The mice were then infected with infected with 400 pfu of PR8 IAV intranasally as depicted in (**C**). Survival of mice **(D**) was tracked daily. n = 9/group. Survival differences were statistically determined by the Gehan-Breslow-Wilcoxon test. Schematics created in BioRender. *p < 0.05

IAV infection upregulates both the EP2 and EP4 receptors on AMs (Coulombe et al., 2014). We examined whether prophylactic blockade of the EP4 receptor would increase survival in aged mice to IAV. Aged mice were given a 7-day prophylactic blockade of EP4 receptor or vehicle control followed by a lethal IAV infection (Supplemental Figure 3A). The prophylactic treatment with the EP4 receptor antagonist failed to increase survival of aged mice to IAV infection (Supplemental Figure 3B, survival tracked up to 15 dpi).

To investigate synergy between the EP2 and EP4 receptor blockades, we administered both the EP2 and EP4 receptor antagonists prophylactically for 7 days pre-infection and to 4 dpi (days -7 to +4) to aged mice and infected the mice intranasally with IAV (Figure 3C). Under this dosing regimen, aged mice treated with the dual EP2/EP4 antagonistic exhibited a significant increase in survival versus those treated with vehicle control aged (55% survival in treated vs. 10% vehicle control, survival tracked up to 16 dpi) (Figure 3D). However, there were no survival differences between the aged mice given the EP2 antagonist treatment alone versus the dual EP2/EP4 receptor antagonists (Supplemental Figure 3C). Thus, dual blockage treatment does not provide additional therapeutic advantages compared to the single EP2 blockade treatment.

In contrast to aged mice, young C57BL/6 mice treated prophylactically with the EP2/EP4 antagonist (Supplemental Figure 3D) and infected with a lethal dose of IAV (i.e., 800 pfu) failed to exhibit improved survival (Supplemental Figure 3E, survival tracked up to 13dpi). These results are compatible with our observation that E2 blockade failed to increase AM numbers prior to infection in young mice (Supplemental Figure 2C). Therefore, excessive PGE_2_ signaling limits both AM numbers and compromises survival to IAV in an age-dependent manner.

We next examined if EP2 blockade could alter the clinical response to IAV in aged mice when the blockade is administered during active infection. Therefore, we infected aged C57BL/6 mice with IAV and administered 7 daily i.p. injections of the EP2 receptor antagonist, or vehicle control, starting on the day of infection (Supplemental Figure 3F). We found no differences in survival between the treated and control mice when administering EP2 blockade at the time of infection (Supplemental Figure 3G, survival tracked up to 16 dpi). IAV infection is known to deplete AMs (Wong et al., 2017). By starting the EP2 blockade on the day of infection, we suspect that AM numbers were not sufficiently boosted during IAV infection to affect survival.

### Prophylactic blockade of PGE_2_ signaling through the EP2 receptor reduces influenza viral load and decreases disease severity in aged mice

Given that prophylactic EP2 receptor blockade increased survival of aged mice following IAV infection, we next examined if EP2 antagonism affected viral load and lung damage. To monitor viral load, we treated aged C57BL/6 mice prophylactically with the EP2 antagonist or vehicle for 7 days, infected them with IAV, and measured the viral protein hemagglutinin (HA) in lung homogenates collected at 4 dpi. EP2 antagonism significantly reduced viral HA protein levels by ∼3-fold compared to the vehicle control (Figure 4A). The reduction in viral protein was not accompanied by altered levels of the anti-viral type I interferon (IFN)-β or the type II IFN-γ in the lung homogenates (Supplemental Figure 4 A-B). EP2 blockade not only reduced viral load but also lung damage, inferred from a ∼2.5-fold reduced level albumin in the BALF, a marker of lung damage, at 4 dpi as compared to control (Figure 4B). BALF from 4 dpi aged mice treated with the EP2 antagonist also exhibited ∼2-fold reductions in the levels of the proinflammatory cytokines IL-6 and TNF-α (Figure 4C-D). Mice begin to recover from infection and resolve the inflammation in the lungs at 9 dpi, and we found that the EP2 antagonist-treated mice exhibited a ∼1.75-fold increase in the immunosuppressive cytokine IL-10 at 9 dpi (Figure 4E), indicating that the EP2 antagonist-treated mice are better able to resolve the inflammation within the lungs than vehicle-treated counterparts. Overall, these data indicated that the EP2 antagonist treatment prior to IAV in aged mice leads to reduced viral load, inflammatory cytokines, and subsequent lung damage following infection.

**Figure 4:**
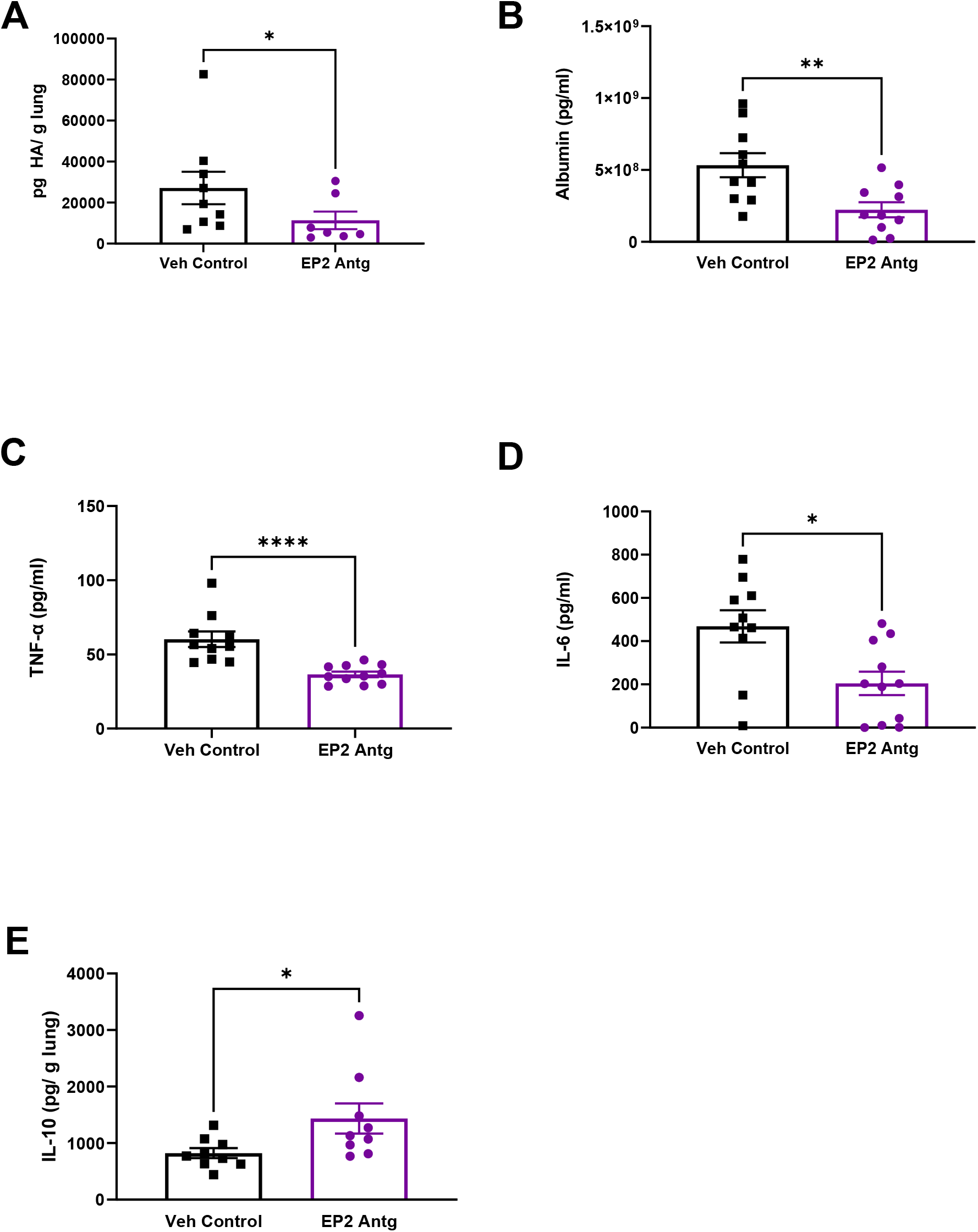
Prophylactic blockade of PGE_2_ signaling through the EP2 receptor reduces influenza viral load and disease severity in aged mice. Aged female C57BL/6 mice were given EP2 antagonist or vehicle control by daily i.p. injection for seven days followed by intranasal (i.n.) infection of PR8 H1N1 as shown in Figure 3A. Lung homogenate and BALF samples were collected on 4 and 9 dpi. **(A)** PR8 H1N1 IAV protein hemagglutinin (HA) measured in the lung homogenate on 4 dpi **(B)** Albumin measured in the BALF on 4 dpi **(C)** TNF-α measured from the BALF on 4dpi **(D)** IL-6 measured from the BALF on 4 dpi (E) IL-10 measured in the lung homogenate on 4 dpi. Statistical significance analyzed by Mann-Whitney. Error bars represent SEM. Each point represents a biological replicate. * p< 0.05, ** p< 0.005, ****p < 0.0001.

As PGE_2_ is a pleotropic lipid known to regulate several immune cell types (Agard et al., 2013; Sander et al., 2017), we characterized how prophylactic blockade of the EP2 receptor affects immune cell accumulation into the lung and BALF following IAV infection. We prophylactically treated aged mice with the EP2 antagonist or vehicle, subsequently infected with IAV, and then performed flow cytometry on the lungs and BALF collected on 9 dpi (gating strategy shown in Supplemental Figure 4C). While total CD45^+^ hematopoietic cell numbers in the BALF were similar between the EP2 antagonist treated and vehicle control groups (Supplemental Figure 4D), there were increased percentages of neutrophils (CD45^+^ Ly6G^+^ CD11b^+^) (Supplemental Figure 4E) and AMs (CD45^+^ SiglecF^+^ CD11c^+^) (Supplemental Figure 4F) within the CD45^+^ population of the BALF in the EP2 antagonist treated mice as compared to controls. There were no differences in total CD45^+^ hematopoietic cell numbers (Supplemental Figure 4G), percentage of CD4^+^ T cells (CD3^+^ CD4^+^CD8^-^) (Supplemental Figure 4H) within the total T cell population, B cells (CD45^+^ B220^+^) (Supplemental Figure 4I) within the CD45^+^ population, or total monocytes/macrophages (CD45^+^ F4/80^+^) (Supplemental Figure 4J) within the lung.

We also used flow cytometry to determine the total number of type II alveolar epithelial cells (AECs) (CD45^-^ EpCAM^+^) of IAV infected aged mice given the prophylactic EP2 antagonist treatment or vehicle control at 9 dpi. Type II AECs contribute to epithelial regeneration following tissue damage (Crosby and Waters, 2010; Zemans et al., 2011) and are a primary cellular target of IAV (Villalón-Letelier et al., 2017; Weinheimer et al., 2012). Interestingly, EP2 antagonist treated aged mice exhibited statistically higher cell counts of type II alveolar epithelial cells AECs compared to the vehicle control (Supplemental Figure 4K). These results suggest that EP2 antagonist treated mice experience less loss of type II AECs during IAV infection compared to vehicle control mice.

### PGE_2_ signaling alters the mitochondrial fitness of AMs

Reduced mitochondrial function, specifically the inhibition of the electron transport chain and oxidative phosphorylation, has been shown to limit proliferation in a variety of cell types such as intestinal stem cells (Zhang et al., 2020), Jurkat cells (Birsoy et al., 2015), vascular smooth muscle cells (Chiong et al., 2014), and human colon cancer cells HCT116 (Wheaton et al., 2014). Additionally, PGE_2_ has been shown to limit oxidative phosphorylation in human monocyte-derived macrophages (Minhas et al., 2021). Hence, we hypothesized that PGE_2_ reduces the mitochondrial fitness and energetic output of AMs, which may inhibit AM proliferation. To test this, we gave aged mice 7 daily i.p., injections of the EP2 antagonist or vehicle control and then collected AMs via BAL. The AMs were analyzed for mitochondrial mass via MitoTracker, mitochondrial reactive oxygen species (ROS) via MitoSOX, and mitochondrial membrane potential via tetramethylrhodamine methyl ester (TMRM) staining. Our results indicate that blocking the EP2 receptor reduced mitochondrial mass (Figure 5A), mitochondrial ROS (Figure 5B), and mitochondrial membrane potential (Fig 5C) of AMs. Additionally, we complemented these findings via *ex vivo* culture of AMs, isolated from young C57BL/6 mice, that were cultured overnight with 0 or 1µM of PGE_2_. Subsequent staining and analysis of cultured AMs revealed that PGE_2_ increases MitoTracker staining for mitochondrial mass by ∼1.3-fold (Figure 5D), providing further evidence that PGE_2_ increases mitochondrial mass of AMs.

**Figure 5:**
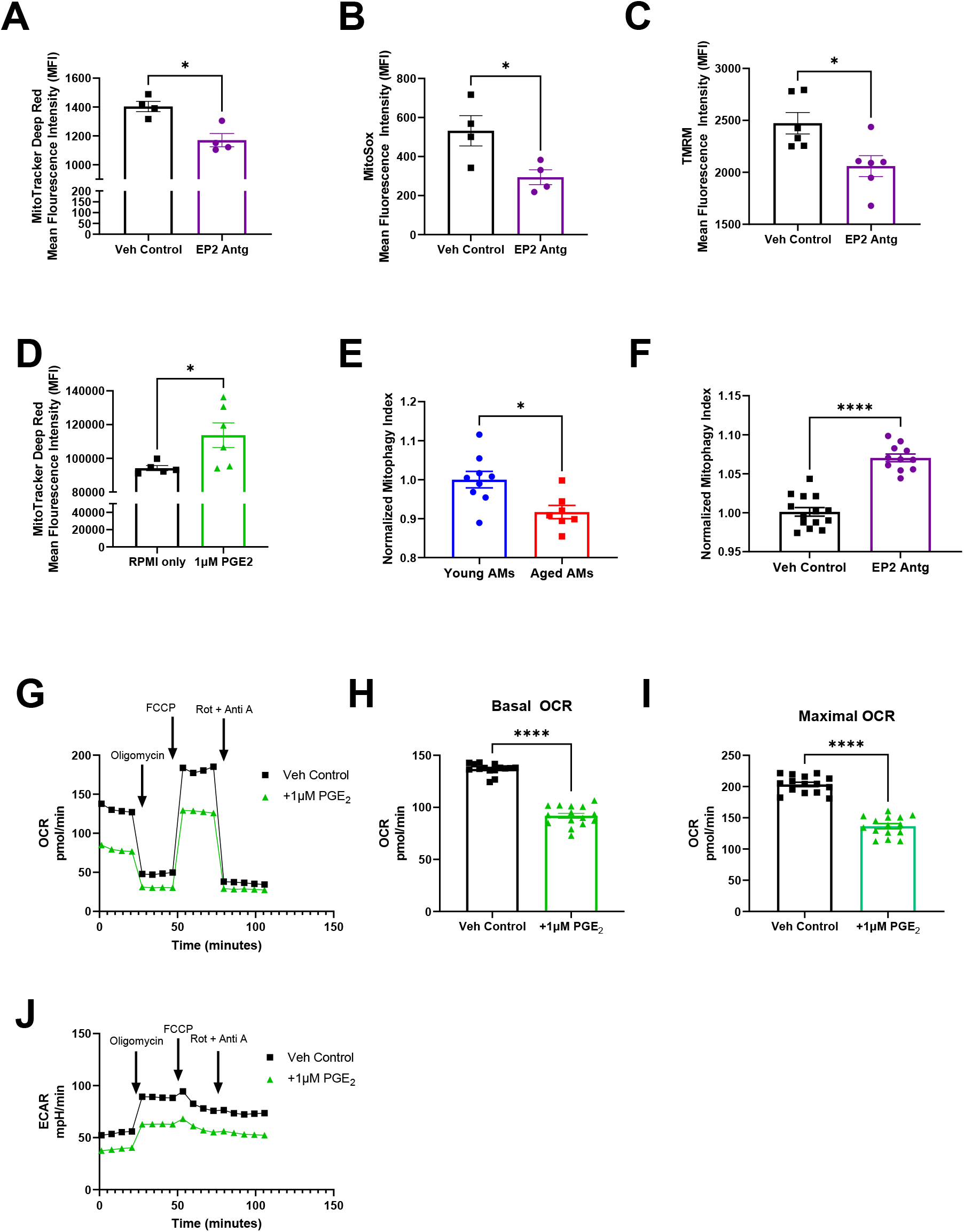
PGE_2_ signaling alters the mitochondrial fitness of AMs. (**A-C**) Aged C57BL/6 mice were given daily injections of either EP2 antagonist or vehicle control by i.p. for 7 days. AMs (i.e., CD45^+^ CD11c^+^ SiglecF^+^) were then collected through BAL, stained with (**A**) MitoTracker DeepRed, (**B**) MitoSox, or (**C**) TMRM, and then analyzed by flow cytometry. The mean fluorescence intensity (MFI) was quantified. (**D**) Primary isolated AMs isolated from young mice were cultured with 1µM of PGE_2_ for 24 hours. Cells were then stained with MitoTracker DeepRed and then analyzed by flow cytometry. The MFI was quantified. (**E-F**) AMs (i.e., SiglecF^+^) were collected through BAL of MitoQC mice and analyzed by flow cytometry. Mitophagy index was calculated based on the mCherry and GFP signals. Mitophagy indexes were normalized within experiments to the control (i.e., young mice or vehicle control treated mice). (**E**) AMs from young (2-3 months) and aged (22-25 months) non-infected MitoQC mice. (**F**) AMs from non-infected MitoQC (9-10 months) mice, given daily injections of either EP2 antagonist or vehicle control by i.p. for 7 days (**G-J**) MH-S cells were cultured for 24 hours with 1µM PGE_2_ or vehicle control. OCR and ECAR were analyzed by a Seahorse xFe96 analyzer. n=15 / group (**G**) OCR measurements (**H**) Basal OCR measurements (**I**) Maximal OCR measurements (**J**) ECAR measurements Statistical significance analyzed by Mann-Whitney. Error bars represent SEM. Each point represents a replicate. * p< 0.05, ** p< 0.005, ****p < 0.0001.

A possible explanation for PGE_2_ increasing mitochondrial mass on AMs is via decreased mitophagy. Mitophagy is the autophagic recycling of damaged mitochondrial (Palikaras et al., 2018), and is dysregulated with aging (Chen et al., 2020a). However, how aging affects mitophagy in AMs is unknown. To determine if aging alters mitophagy of AMs, we utilized a mitophagy reporter mouse model, the MitoQC mice. MitoQC mice contain a pH-sensitive GFP and mCherry mitochondrial marker. Under neutral pH, both the mCherry and GFP fluoresce, however within the acidic environment of the autolysosome the GFP fluorescence is quenched. The MitoQC mice allow for the *in vivo* detection of mitophagy at a single-cell level by measuring the GFP and mCherry fluorescence signals (McWilliams et al., 2016). AMs were obtained from the BALF of non-infected young (2-3 months) and aged (22-25 months) MitoQC mice. We then gated on the AM population (i.e, SiglecF^+^) and analyzed the mCherry and GFP fluorescence signals via flow cytometry. The mitophagy index was then calculated based on the mean fluorescence intensity (MFI) of mCherry divided by MFI of GFP, and the results normalized to control conditions (i.e., AMs from young mice, or AMs from vehicle control treated mice). We found that when AMs from aged MitoQc mice were compared to AMs from young MitoQC mice, aged AMs exhibited a reduced mitophagy index (Figure 5E). We next determine if PGE_2_ is an age-associated factor that limits mitophagy in AMs, by treating middle-aged (i.e., 9-10 months) non-infected MitoQC mice with seven daily doses of the EP2 antagonist. When we compared EP2 antagonist treated mice to vehicle control, we found that blocking the EP2 receptor increased mitophagy in AMs *in vivo* (Figure 5F). Overall, results indicate that aging, via elevated PGE_2_, restricts mitophagy in AMs, potentially leading to an accumulation of damaged mitochondrial.

To understand how PGE_2_ affects cellular metabolism in AMs, we measured the oxygen consumption rate (OCR), a readout of oxidative phosphorylation, and extracellular acidification rate (ECAR), a readout of glycolysis, in AMs cultured with PGE_2_ via a Seahorse assay (Agilent) (Zhang et al., 2012). As it was not practical to isolate sufficient numbers of primary AMs for this assay, we employed an immortalized murine AM cell line, MH-S (Mbawuike and Herscowitz, 1989; Sankaran and Herscowitz, 1995). We first noted that MH-S cells increased mitochondrial ROS (Supplemental Figure 5A-B), increased mitochondrial membrane potential (Supplemental Figure 5C-D), and reduced proliferation when cultured with PGE_2_ (Supplemental Figure 5E). These findings are compatible with our *in vivo* results in which EP2 antagonism decreased mitochondrial ROS, decreased mitochondrial membrane potential in AMs (Figure 5B-C), and increased AM cell numbers (Figure 2A). Given this, we employed the MH-S cell line in the Seahorse assay, which revealed that PGE_2_ lowered OCR over the course of the assay (Figure 5G), including reductions in both basal OCR (Figure 5H), and maximal OCR (Figure 5I) by ∼1.5-fold. Additionally, PGE_2_ limited the ECAR of MH-S cells (Figure 5J). Overall, these results suggest that PGE_2_ restricts both oxidative phosphorylation and glycolysis of AMs, to impair mitochondrial homeostasis and energy generation in AMs, and ultimately reduce AM proliferation.

### Senescent type II AECs are a primary source of PGE_2_ in the aged lung

Next, we determined which cell type is a major contributor to the elevation of PGE_2_ in the BALF with aging. At steady state, AMs are the predominant cell type in the lung airways and airspace. Macrophage populations, such as AMs, peritoneal macrophages and microglial cells, are known producers of PGE_2_ (Balter et al., 1988; Minhas et al., 2021; Williams and Shacter, 1997). Type I and type II alveolar epithelial cells line the alveoli and secrete factors into the airspace. Type I AECs provide structure and barrier defense whereas type II AEC are highly secretory cells and are readily characterized by their lipid-rich lamellar bodies (Whitsett and Alenghat, 2015). We re-analyzed publicly available scRNA-seq data of AECs (GEO GSE113049) and found that type II AECs express ∼5.5-fold higher transcript levels of COX-1 and ∼8.5-fold higher transcript levels of COX-2, critical enzymes for the rate-limiting step of PGE_2_ synthesis, relative to type I AECs (Supplemental Figure 6A) (Herschman and Hall, 1994; Riemondy et al., 2019). Therefore, we considered AMs and type II AECs as two main candidate cell types responsible for PGE_2_ in the BALF.

To determine whether AMs or type II AECs are a predominant source of PGE_2_ with aging, we isolated and cultured both AMs and type II AECs from non-infected young and aged mice. Following 2 days of culture, the cell culture medium was collected and analyzed for PGE_2_ by ELISA. Type II AECs produced PGE_2_ at 3 orders of magnitude higher than AMs (Figure 6A). Importantly, type II AECs from aged mice produced about 2-fold more PGE_2_ than type II AECs from young mice (Figure 6A). These results suggest that type II AECs, not AMs, are a major source of PGE_2_ in the BALF before infection, and aging increases the production of PGE_2_ by type II AECs.

**Figure 6:**
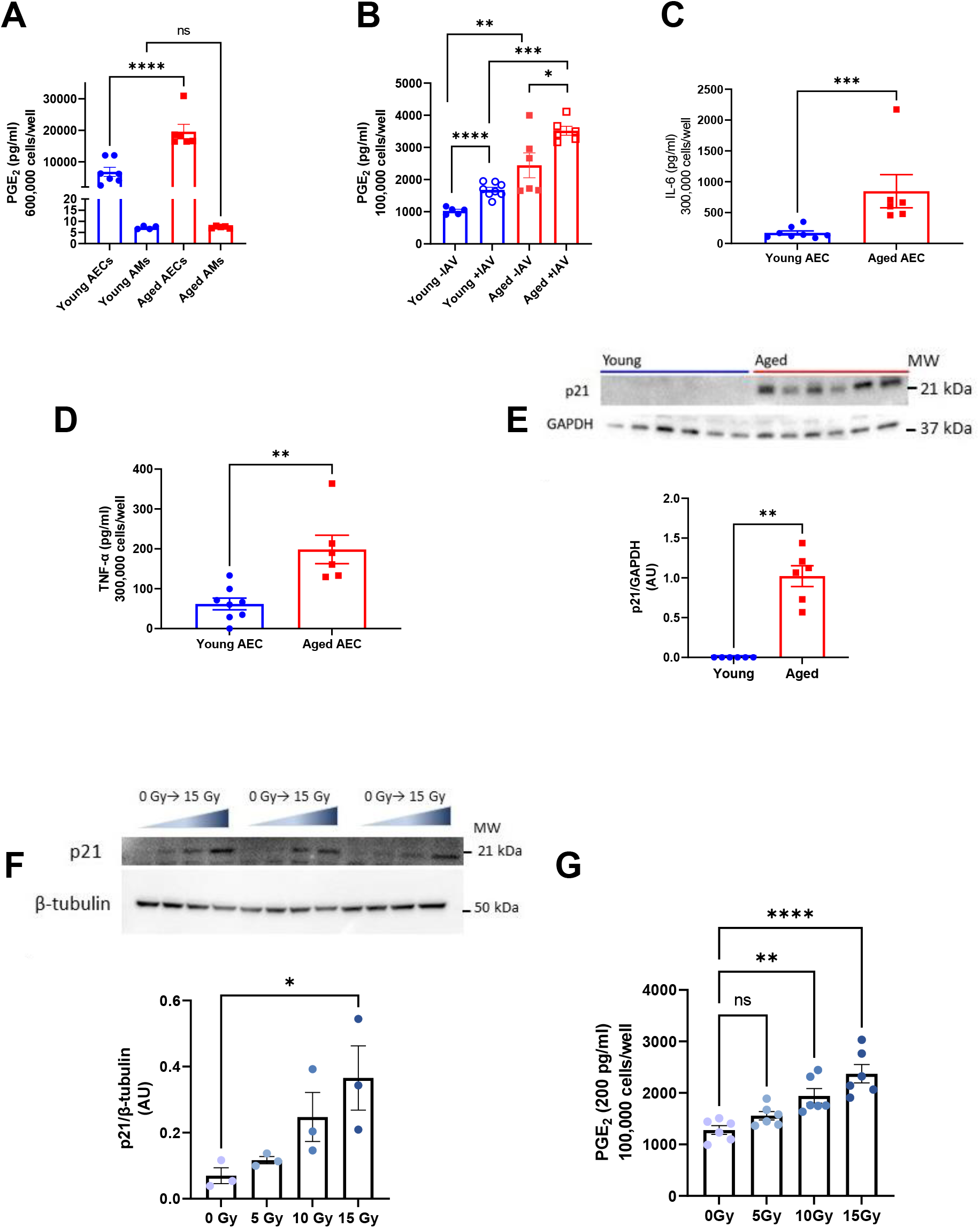
Senescent type II AECs are a primary source of PGE_2_ in the aged lung. (**A-D**) Primary type II AECs and AMs were isolated from young and aged C57BL/6 mice and then cultured *ex vivo*. Cell culture media following 2 days of accumulation was collected and analyzed by ELISA for secreted factors. (**A**) Basal PGE_2_ secretion by primary type II AECs and AMs in culture. Statistical significance analyzed by ANOVA with Tukey post-hoc test. (**B**) Primary type II AECs were isolated from young and aged C57BL/6 mice and were infected *ex-vivo* with PR8 IAV (MOI = 0.1). Cell culture media was collected following 2 days of accumulation and analyzed for PGE_2_ with ELISA. Statistical significance analyzed by ANOVA with Tukey post-hoc test (**C**) IL-6 secreted by primary type II AECs in culture. Statistical significance analyzed by Mann-Whitney. (**D**) TNF-α secreted by primary type II AECs in culture. Statistical significance analyzed by Mann-Whitney. (**E**) Primary type II AECs isolated from young and aged C57BL/6 mice were lysed and analyzed for p21 by Western blot. Statistical significance analyzed by Mann-Whitney. (**F-G**) Primary type II AECs isolated from young C57BL/6 mice were irradiated with increasing levels of radiation (i.e., 0, 5, 10, 15 Gy) to induce senescence. (**F**) Expression of p21 was quantified and normalized based on β-tubulin. Statistical significance analyzed by ANOVA with Tukey post-hoc test. (**G**) Cell culture media was collected and analyzed for PGE_2_ levels by ELISA. Statistical significance analyzed by ANOVA with Tukey post-hoc test. Error bars represent SEM. Each point represents a biological replicate. *p<0.05, **p<0.005, ***p<0.0005, ****p<0.0001.

We next determined if IAV infection increases PGE_2_ production by type II AECs and whether aging exacerbates the phenotype. Hence, we cultured type II AECs isolated from either young or aged mice and infected the AECs *in vitro* with IAV at a multiplicity of infection (MOI) of 0.1 for 2 days. We subsequently measured PGE_2_ in the culture supernatants via ELISA. Importantly, IAV led to increased PGE_2_ production by ∼1.5-fold in both young and aged AECs, but infected type II AECs from aged mice still produced ∼2-fold higher levels of PGE_2_ as compared to infected type II AECs isolated from young mice (Figure 6B).

### Senescence is sufficient to increase PGE_2_ production by type II AECs

Next, we sought to determine a cellular mechanism by which aging increases PGE_2_ production by type II AECs. One of the hallmarks of aging is cellular senescence, a phenomenon characterized by the irreversible arrest of the cell cycle (Kumari and Jat, 2021). Senescent cells accumulate with age and secrete pro-inflammatory factors at steady state through a phenomenon known as the SASP (Di Micco et al., 2021; Kumari and Jat, 2021). To determine if senescence increases PGE_2_ secretion by type II AECs and if PGE_2_ is a SASP factor, we first characterized whether type II AECs from aged mice exhibit features of senescence. Hence, we measured the secretion of two SASP factors, IL-6 and TNF-α from type II AECs isolated from non-infected young and aged mice (Figure 6C-D). Notably, type II AECs from non-infected aged mice exhibited a ∼2-3 fold increased secretion of both IL-6 and TNF-α as compared to type II AECs from non-infected young mice. Additionally, we measured the senescence marker p21 in young and aged type II AEC lysates by western blot (Figure 6E). Type II AECs from non-infected young mice had undetectable levels of p21, whereas type II AECs from non-infected aged mice exhibited marked expression of p21. Overall, these results show that type II AECs isolated from non-infected aged mice exhibit evidence of senescence.

To determine if senescence is sufficient to increase PGE_2_ secretion, we irradiated type II AECs isolated from young mice to induce a senescent phenotype within the cells (Li et al., 2018). The type II AECs received either 0Gy, 5Gy, 10Gy, or 15Gy of radiation. We then measured p21 by western blotting the irradiated cells to confirm the induction of a senescence phenotype. The levels of p21 in type II AECs increased with increasing levels of irradiation (Figure 6F). We then measured PGE_2_ in the cell culture medium of the irradiated type II AECs by ELISA. With increasing levels of radiation, there was increased PGE_2_ production (Figure 6G). Additionally, our results show a positive correlation between p21 expression and PGE_2_ production in irradiated type II AECS (Supplemental Figure 6B). Together, these results suggest that PGE_2_ is a SASP factor of senescent type II AECs, and that senescent type II AECs are a major producer of PGE_2_ with aging.

## Discussion

The dysregulation of the immune system with age, called immunosenescence, contributes to increased morbidity and mortality to respiratory pathogens, including IAV (Chen et al., 2020b). While the effects of immunosenescence on adaptive immunity are well-studied, the effects of immunosenescence on the innate immune system are less well understood (Aw et al., 2007; Chen et al., 2020b). Here, we show that the overproduction of PGE_2_ contributes to the immunosenescence of AMs. We show that aged mice exhibit increased levels of the lipid PGE_2_ both systemically in the plasma and in the BALF. Prophylactically blocking PGE_2_ signaling increases the number of AMs prior to infection and subsequently enhances the survival of aged, but not young, mice to lethal IAV infection. Thus, age-elevated PGE_2_ is detrimental to host immunity against IAV infection.

AMs are tissue resident macrophages of the airways and airspaces that have established functions in tissue homeostasis, host defense, and resolving inflammation (Hussell and Bell, 2014). We have previously shown that higher PGE_2_ levels in the BALF is correlated to reduced AM numbers *in vivo*, and that PGE_2_ restricts AM proliferation *in vitro* (Penke et al., 2020). These findings suggest that PGE_2_ reduces AMs numbers, a finding that is compatible with our prior work in which we found that AMs numbers decrease with aging (Penke et al., 2020; Wong et al., 2017). We, as well as others, have shown the importance of AMs to lethal IAV infections through the genetic or pharmacological depletion of AMs (Cardani et al., 2017; Wong et al., 2017). Our study demonstrates that blocking PGE_2_ function via inhibiting the EP2 receptor is sufficient to increase AM numbers via increased proliferation in non-infected aged mice. Our study, however, does not exclude the possibility of increased recruitment of monocytes into the airways and their differentiation into AMs. However, since monocytes are typically recruited to the lung under inflammatory conditions rather than at steady state, we do not suspect that monocyte recruitment is a major contributor to the increased AMs identified prior to infection in our model (Goto et al., 2004; Sen et al., 2016).

Our study suggests that increasing AM numbers with aging, via EP2 receptor blockade, augments innate immune defense and inflammation resolution during IAV. This could explain several observations of our study, including that the EP2 receptor blockade in aged mice reduced viral burden, reduced inflammatory cytokines TNF-α and IL-6, and importantly increased survival to lethal IAV infection with aging. Additionally, this could explain the observation of higher number of type II AECs, the primary cellular target of IAV, in the lung at 9 dpi in the EP2 antagonist treated mice (Villalón-Letelier et al., 2017; Weinheimer et al., 2012). We hypothesize that the improved AM-mediated innate immunity and control of IAV in the EP2 antagonist treated mice lead to reduced type II AEC death. This is further supported by the reduction of the albumin in the BAL, a marker of lung damage. Overall, our data suggest that the prophylactic EP2 receptor blockage improves host defense against IAV with aging.

We also found that whereas EP2 receptor blockade increased AM numbers in non-infected aged mice, it failed to do so in non-infected young mice. It is possible that in young mice, who have lower PGE_2_ levels relative to aged mice, PGE_2_ levels are already optimized so that EP2 blockade is ineffectual at increasing AM numbers. This might explain why blocking PGE_2_ signaling only improved survival to IAV in aged, but not young, mice. Finally, it is likely that 7 days of prophylactic treatment of EP2 blockade sufficiently boosted the number of AMs to improve outcomes after IAV infection in aged mice, whereas starting EP2 blockade on the day of infection likely did not give sufficient time to increase AM numbers during IAV infection, which is known to deplete AMs. This may explain why starting treatment with EP2 blockade at the time of infection failed to increase survival in aged mice following IAV infection.

Our study provides a mechanistic basis for why age-related elevations of PGE_2_ lead to lower numbers of AMs. Specifically, we provide evidence that PGE_2_ impedes proliferation of AMs. As AMs are cells that self-renew within the lung, our results suggest that PGE_2_ restriction of AM proliferation likely impairs self-renewal of AMs within the aged lung. Our study also indicates that PGE_2_ perturbs the metabolic health of AMs via mitochondrial dysfunction, a hallmark of aging (López-Otín et al., 2013). Importantly, defective or damaged mitochondria are removed from the cell via a process termed mitophagy, a specific form of macro-autophagy (Palikaras et al., 2018). Altered mitophagy, which occurs with aging, contributes to age-associated diseases including atherosclerosis, Parkinson’s disease and chronic lung disease (Chen et al., 2020a; Ito et al., 2015; Liu et al., 2019; Sureshbabu and Bhandari, 2013; Tyrrell et al., 2020). Employing a mitophagy reporter mice, our study has found that activation of the EP2 receptor, the major PGE_2_ receptor on AMs, limits mitophagy in AMs. Our study found that EP2 receptor blockade reduces mitochondrial mass, mitochondrial oxidative stress and mitochondrial membrane potential in AMs, which could be explained by EP2 receptor blockade increasing mitophagy. Our study also shows that PGE_2_ restricts both oxidative phosphorylation and glycolysis of AMs. However, it is possible that PGE_2_ impacts mitochondrial health in AMs independently of its effects on mitophagy. Whether PGE_2_ restricts mitochondrial health, via mitophagy or other mechanisms, in other innate immune cells and other immune cells in general, and the possible physiological consequences thereof, will require future investigation.

We identified senescent type II AECs as a predominant cellular source of elevated PGE_2_ in the aged lung. Our study suggests that with aging, type II AECs communicate aberrantly with AMs via the excessive PGE_2_ secretion, which impairs AM proliferation as stated above. Interestingly, PGE_2_ has been shown to induce and maintain senescence in human fibroblasts and CD8^+^ T cells (Chou et al., 2014; Martien et al., 2013). Whether PGE_2_ participates in a positive-feedback loop with the senescence phenotype in AECs, or other cells in the lung, has not yet been studied. We speculate that the increased PGE_2_ levels within the lung with aging further promotes a senescence phenotype in the AECs, leading to sustained production of PGE_2_. However, this speculation requires future investigation.

It is unlikely that the role of PGE_2_ in immunosenescence uncovered here is specific to IAV infection. The applications of our study may be expanded to other respiratory infections such as COVID-19. Aging is a prominent risk factor for severe COVID-19 and COVID-19 related mortality. Interestingly, COVID-19 disease severity is positively correlated with PGE_2_ serum levels (Ricke-Hoch et al., 2021). Additionally, infecting Calu-3 cells, a human lung epithelial cell line, with SARS-CoV-2 induced PGE_2_ production (Ricke-Hoch et al., 2021); similar to our finding that IAV infection increases PGE_2_ levels in the lungs *in vivo,* and promotes PGE_2_ production by primary type II AECs *ex vivo*. Besides respiratory viruses, AMs also play critical roles in the immune defense against respiratory bacteria such as *Streptococcus pneumoniae,* and *Staphylococcus aureus* (Ghoneim et al., 2013; Knapp et al., 2003; Yajjala et al., 2016). The limiting of AM cell numbers by increased PGE_2_ in the lungs of older adults may play a role in the high incidence of pneumonia and pneumonia-related morbidity in older adults (Childs et al., 2019; Ebright and Rytel, 1980). In addition to proliferation, PGE_2_ has been shown to limit AM phagocytosis, killing of bacterial cells and the expression of the toll-like receptor 4 (Aronoff et al., 2004; Degraaf et al., 2014). This may represent another mechanism by which age-enhanced PGE_2_ levels in the lungs limit AM-mediated immunity. The role of the age-enhanced PGE_2_ levels in the lungs in regulating AMs and other immune cells and its implications in other respiratory diseases will require future investigation.

## Conclusion

In conclusion, our study has revealed that age-associated overproduction of PGE_2_ in the lung, largely by senescent type II alveolar epithelial cells, impairs AM proliferation, and reduces total AM cell numbers. This limits the ability of AMs to defend against IAV infection. Additionally, our study shows that PGE_2_ impairs mitophagy and mitochondrial health of AMs (graphically represented in Figure 7). Thus, we have identified a novel, aberrant form of cross talk between type II AECs and AMs, via the secretion of PGE_2_, that restrains AM numbers to compromise host defense to IAV infection with aging. Additionally, our study suggests that potential therapies that target senescent type II AECs and their production of PGE_2_ or the EP2-mediated signaling in AMs, may reduce the burden of IAV, and possibly other respiratory viruses such as coronaviruses, in older adults.

**Figure 7:**
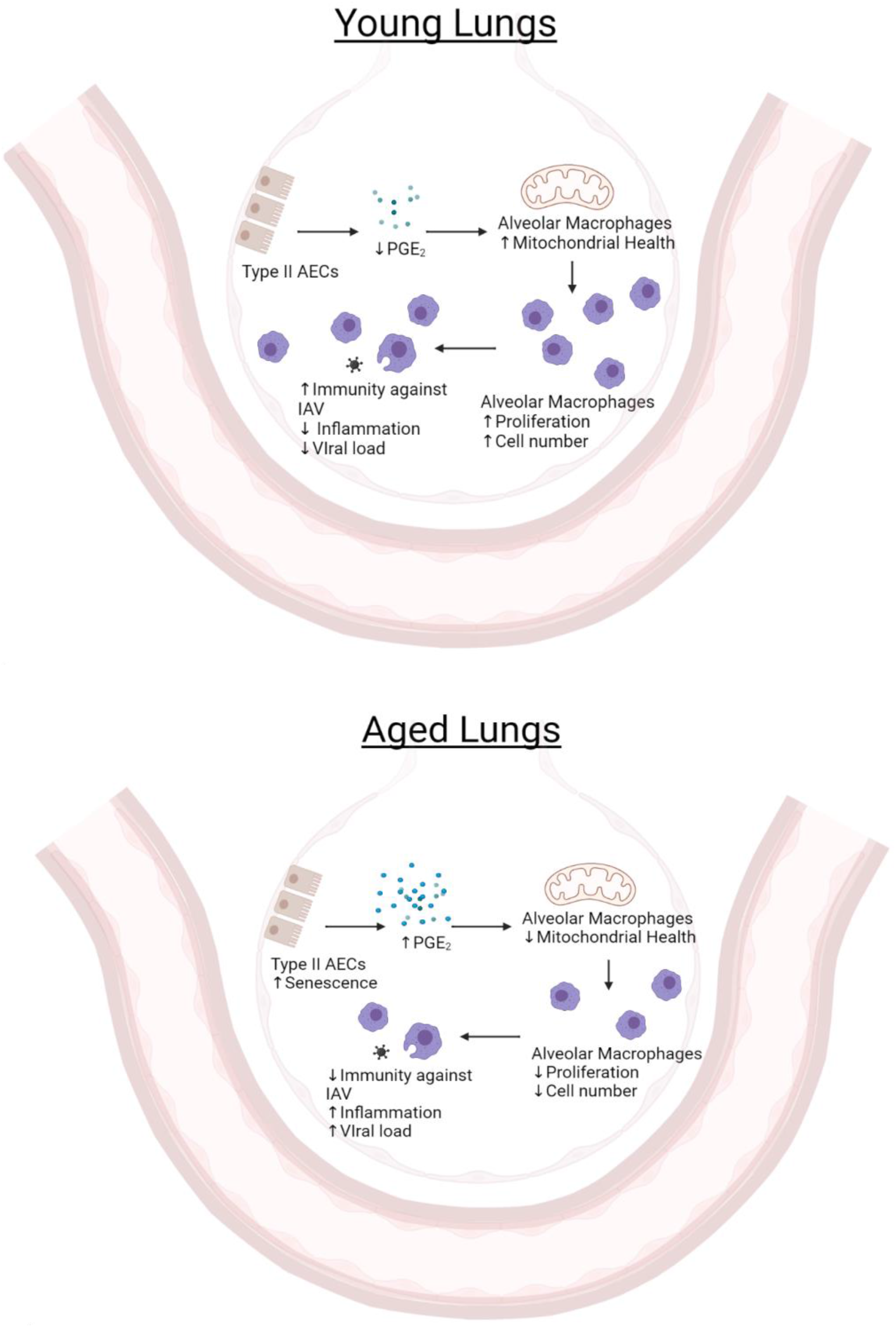
Aging leads to pathological communication between AECs and AMs via PGE_2_ secretion. **Top:** With youth there are low secreted concentrations of PGE_2_ by AECs which do not impact AM mitochondrial function. Hence, there are sufficient AM numbers to maintain homeostasis and provide host defense to influenza infection. **Bottom**: With aging, senescent AECs secrete increased levels of PGE_2_ that subsequently limit AM mitochondrial function, proliferation and reduce AM numbers. Consequently, host defense to influenza viral infection is compromised. Schematics created in BioRender.

## Experimental Procedures

See the Supplemental Information for details.

### Study approval

All animal experiments were approved prior to the initiation of the study and were carried out in accordance with the University of Michigan Institutional Animal Care and Use Committee.

### Mice

Young (2-4 months) and aged (18-22 months) female C57BL/6 mice were obtained from Charles Rivers and the National Institute of Aging rodent facility at Charles Rivers. Male UM-HET3 mice were kindly gifted by Dr. Richard Miller at the University of Michigan. The UM-HET3 mice were aged at the Glenn Center on Aging at the University of Michigan. MitoQC mice were donated from the lab of Dr. Ian Ganley at the University of Dundee and originally generated by Taconic Artemis (McWilliams et al., 2016). The MitoQC mice were then bred and housed within the animal facility at the North Campus Research Complex at the University of Michigan. All mice were maintained on a 12-hour light-dark cycle with free access to food and water within a specific-pathogen-free facility. Mice were monitored for at least 1 week after arrival to our facilities for signs of stress and/or disease. Animals that displayed evidence of infection or illness prior to influenza infection were excluded from the study.

### PGE_2_ receptor antagonists *in vivo*

The EP2 antagonist PF-04418948 (catalog # 15016) and the EP4 antagonist ONO-AE3-208 (catalog # 14522) were purchased from Cayman Chemical. Mice were given daily doses of 10mg/kg EP2 and/or 10mg/kg EP4 antagonist(s) by intraperitoneal (i.p.) injections

### Virus

Stocks of the VR-95 stain of Influenza A/PR8/34 H1N1 were purchased from ATCC.

### *in vivo* influenza infection

Mice were anesthetized with isoflurane and instilled with 400 plaque forming units (PFU) of influenza A virus in 40µl PBS or 40µl PBS vehicle control.

### *in vitro* influenza infection

Cells were infected with an MOI= 0.1 of influenza virus diluted in cell culture medium for 48 hours.

### Flow Cytometry

Cells were obtained from BAL, single-cell suspension of lungs or cell culture. Flow cytometry was performed using the ZE5 Cell Analyzer (BioRad) of the Flow Cytometry Core at the University of Michigan. Analysis of flow cytometry data was performed using FlowJo (version 10.8.0).

### MH-S cell culture

MH-S cells, an immortalized mouse alveolar macrophage cell line (Mbawuike and Herscowitz, 1989), were grown in RPMI-1640 medium containing 10% FBS, 0.05mM 2-mercaptoethanol, and 100U/ml penicillin/streptomycin. For PGE_2_ stimulation assays, cells were cultured with 1 µM PGE_2_, 10 µM PGE2, or vehicle control, for 24 hours.

### Isolation and culture of type II AECs

AECs were isolated by magnetic associated cell sorting (MACS) by negative selection of CD45^+^ and CD31^+^ cells and followed by positive selection of CD326^+^ cells.

Isolated type II AECs were resuspended in a DMEM/F12 medium containing 10% FBS, 1.25g BSA, 100U/mL penicillin/streptomycin, and 1x Insulin-Transferrin-Selenium (Gibco, 41400045). Type II AECs were seeded in tissue culture plates coated with gelatin-based coating solution (Cell Biologics, 6950).

### Seahorse Assay

Four basal readings were taken prior to the addition of the electron transport chain inhibitors in Agilent’s mitostress test: 1.5µM oligomycin, 1µM FCCP, and 0.5µM rotenone and antimycin A (Agilent, 103708). OCR and ECAR data were normalized to total cellular protein levels as measured by BCA assay (ThermoFisher, 23225). Seahorse experiments were repeated three times.

### BrdU in vivo

BrdU (Biogems, 5911439) was given to mice in their drinking water *ad libitum* for a total of 7 days. BrdU was dissolved in the drinking water at a concentration of 0.8mg/ml and the BrdU water was refreshed every 48 hours.

### Statistics

All results are presented as mean ± standard error of the mean (SEM). Data was analyzed using the nonparametric Mann-Whitney unless otherwise stated. Multiple comparison testing was done by ANOVA with Tukey post hoc. Survival differences were analyzed using a Gehan-Breslow-Wilcoxon test. Specific statistical tests are denoted in the figure legends. Two-sided p-values < 0.05 were considered significant. Prism 9 (GraphPad) was used for statistical testing and the generation of graphs.

## Acknowledgements

This work was supported by the NIA AG028082 and NHLBI R35 HL155169 awarded to DRG; the T32 AI 007413 awarded to the Program in Immunology at the University of Michigan; the NHLBI F31 HL158003 awarded to JC; the Veterans Administration award 5 I01 BX004565 awarded to JCD; and the NHLBI R35 HL144979 awarded to MPG. Seahorse experiments were conducted at the Adipose Tissue Core of the Michigan Nutrition Obesity Research Center supported by P30 DK089503.

The authors would like to thank Dr. Richard Miller at the University of Michigan for donating the UM-HET3 mice. We would also like to thank Dr. Peng Jiang for performing preliminary technical experiments with AECs.

## Author Contributions

JC: Conceptualization, data collection, methodology, data analysis, wrote original draft of manuscript, edited manuscript. JCD: Conceptualization, edited manuscript. RZ: Conceptualization, edited manuscript, data analysis. MZ: Conceptualization, edited manuscript. MPG: Conceptualization, edited manuscript. DRG: Conceptualization, supervision, funding acquisition, access to data, wrote and edited manuscript

