## Supplemental Information for "Age-elevated prostaglandin E_2_ enhances mortality to influenza infection"

### Supplemental Experimental Procedures

#### $PGE_2$ receptor antagonists *in vivo*

The EP2 antagonist PF-04418948 (catalog # 15016) and the EP4 antagonist ONO-AE3-208 (catalog # 14522) were purchased from Cayman Chemical. The antagonists were reconstituted in DMSO, aliquoted, and stored at -20°C until use. Antagonists were then diluted with sterile PBS prior to use. DMSO diluted in PBS was used as vehicle control. Mice were given daily doses of 10mg/kg EP2 and/or 10mg/kg EP4 antagonist(s) by intraperitoneal (i.p.) injections using 30G insulin syringes. For experiments treating mice with the EP2 and EP4 antagonists starting on the day of influenza infection, mice were given daily injections on day 0 to day +7 post infection. For experiments treating mice with the EP2 and EP4 antagonists both prior to and during influenza infection, mice were given 11 daily injections on day -7 pre infection to day +4 post infection. For experiments treating mice with the EP2 antagonist prior to influenza infection, mice were given daily injections on day -7 to day 0 pre infection.

#### Virus

Stocks of the VR-95 stain of Influenza A/PR8/34 H1N1 were purchased from ATCC. The stocks were titered using a viral plaque assay. The stocks were then aliquoted and stored at -80°C until use.

#### *in vivo* influenza infection

Mice were anesthetized with isoflurane and instilled with 400 plaque forming units (PFU) of influenza A virus in 40µl PBS or 40µl PBS vehicle control. Following the infection, mice were monitored daily for weight change, clinical score, and mortality. Mice were euthanized after loss of 30% of their pre-infection body weight.

**Sample collection and preparation**

Bronchoalveolar lavage fluid (BALF): The trachea was exposed and cannulated with a blunted 20G IV catheter. A small amount of surgical suture was used around the trachea to tie the catheter in place. The BALF was collected by lavaging the lungs twice with 1mL of cold sterile PBS. BALF was centrifuged at 500 g for 5 minutes and the supernatant was stored at -80°C until analysis. Cells in the BALF were resuspended into FACS buffer (PBS + 2mM EDTA + 4% FBS) for flow cytometry staining or RPMI-1640 medium containing 10% FBS and 100U/mL penicillin/streptomycin for *ex vivo* culture.

Plasma: Following euthanasia, blood was collected from mice using cardiac puncture with a 25G needle. The blood was then promptly mixed with 80μL 0.5M EDTA. The blood samples were then centrifuged at 2200 g for 20 minutes at 4°C. Plasma was then collected from the spun samples, aliquoted, and stored at -80°C until analysis.

Lung homogenate: Lungs were collected and placed in a 2mL microcentrifuge tube containing 5 stainless steel beads (Omni) and 1ml of sterile PBS. The lungs were then homogenized using a TissueLyser II (Qiagen) set at 30Hz for 3 minutes. Following, the samples were centrifuged at 500 g for 5 minutes at 4°C. The supernatant was then collected, aliquoted, and stored at -80°C.

**Flow Cytometry**

Sample prep: Lungs were minced and digested in sterile Hank's Buffered Salt Solution (HBSS) containing 1mg/ml Collagenase D (Roche, 11088866001) and 100 U/ml DNase (Roche, 4716728001) for 30 minutes at 37°C. The lungs were then pressed through a 100 micron cell strainer to obtain single cell suspensions. Red blood cells (RBC) were removed using RBC lysis buffer according to manufacturer's directions (eBioScience, 00-4300-54). Cells were then resuspended in FACS buffer.

Mitochondrial analysis: Live cells were stained with MitoTracker Deep Red RM (ThermoFisher, M22426), tetramethylrhodamine methyl ester (TMRM) (ThermoFisher, T668), and MitoSOX (ThermoFisher, M36008) according to the manufacturer's instructions.

Staining: Cells were stained with a live/dead viability dye according to manufacturer's instructions (ThermoFisher, L34966) for 30 minutes. Cells were washed twice with FACS buffer. All centrifugation steps were performed at 500 g for 5 minutes at 4°C. Next, Fc blocking was performed by incubation with anti-CD16/32 antibody for 20 minutes (Biolegend, 101320). Cells were washed twice with FACS buffer. Following, cells were stained with the desired surface markers for 25 minutes. Cells were washed twice with FACS buffer and fixed in 4% paraformaldehyde for 25 minutes. Cells were washed twice and resuspended in FACS buffer until analysis. Flow cytometry was performed using the ZE5 Cell Analyzer (BioRad) of the Flow Cytometry Core at the University of Michigan. Analysis of flow cytometry data was performed using FlowJo (version 10.8.0). For intracellular staining, fixation/permeabilization of cells were done using CytoFix/CytoPerm (BD, 554714) according to manufacturer's instructions, following surface staining.

Antibodies: APC: BrdU (BD, cat# BD552598), CD45 (Biolegend, clone 30-F11), Ki67 (Biolegend, clone 16A8; APC-eFluor 780: CD8 (eBioScience, 53-6.7); Brilliant Violet 421: CD11b (Biolegend, clone M1/70), CD3 (Biolegend, clone 17A2), EpCam (Biolegend, clone G8.8); Brilliant Violet 605: B220 (Biolegend, clone RA3-6B2), Ly6G (Biolegend, 1A8), CD11c (Biolegend, clone N418); Brilliant Violet 750: SiglecF (BD, clone E5U-2440); Brilliant Violet 785: MHC II (Biolegend, clone M5/114.15.2), FITC: AnnexinV (R&D Systems, cat# 4830-01-K); PE: F4/80 (Biolegend, clone BM8), SiglecF (Biolegend, clone E50-2440); PE-Cy7: CD4 (Invitrogen, clone GR1.5); PE-eFlour 610 (Invitrogen, clone N418).

**Enzyme-linked immunosorbent assay (ELISA)**

TNF-α (88-7324-22), IL-6 (88-7064-88), IFN-γ (88-7314-22) and IL-10 (88-7105-22) ELISA kits were purchased from Invitrogen; albumin ELISA kits were purchased from Abcam (ab108792); cyclic AMP (581001) and $PGE_2$ (514010) ELISA kits were purchased from Cayman Chemical; IFN-β ELISA (42410) kits were purchased from pbl Assay Science; Influenza A/PR/8/1934 hemagglutinin (SEK11684) ELISA kits were purchased from SinoBiological, and $PGE_2$ (MBS266212) ELISA kits were purchased from MyBioSource. All ELISAs were performed according to manufacturer's directions. Each sample was analyzed with at least two dilutions and with two technical replicates per dilution.

**MH-S cell culture**

MH-S cells, an immortalized mouse alveolar macrophage cell line, were grown in RPMI-1640 medium containing 10% FBS, 0.05mM 2-mercaptoethanol, and 100U/ml penicillin/streptomycin. For $PGE_2$ stimulation assays, cells were cultured with 1 μM $PGE_2$, 10 μM PGE2, or vehicle control, for 24 hours.

**Isolation and culture of type II AECs**

AECs were isolated by magnetic associated cell sorting (MACS). First, lungs were perfused with 10ml of phosphate buffered saline (PBS). BAL was performed with 1ml of PBS. Then the lungs were instilled with 3ml of dispase diluted 10-fold (Corning, #354235) and 0.5ml of 1% agarose. Lungs were then collected, minced with sterile scissors, and then digested by incubation with 10x diluted dispase and 0.01% DNase I (Roche, 10104159001) in HBSS for 30 minutes. Single cell suspensions of the lungs were obtained by consecutive passage through 100 micron, 70 micron, and 40 micron cell strainers. RBC were removed using RBC lysis buffer according to manufacturer's directions (eBioScience, 00-4300-54). The suspension was then incubated with CD45 (Miltenyi, 130-052-391) and CD31 microbeads (Miltenyi, 130-097-481) in PBS containing 2mM EDTA and 0.5% BSA. Cells were then washed and centrifuged at 500 g for 5 minutes. Cells were then passed through a MACS column in an OctaMACS magnet. The flow through cells ($CD45^-$ $CD31^-$) were incubated with biotin labeled CD326 (EpCAM) antibody (Miltenyi, clone caa7-9G8). Cells were washed and then incubated with anti-biotin microbeads (Miltenyi, 130-090-485). After washing again, the cells were passed through a MACS column. The flow through was discarded. The attached $CD326^+$ type II AECs were removed from the column by firmly applying a plunger. The purity of the flow through was determined by flow cytometry. Typically, we obtained about 85-90% purity from the isolation.

Isolated type II AECs were resuspended in a DMEM/F12 medium containing 10% FBS, 1.25g BSA, 100U/mL penicillin/streptomycin, and 1x Insulin-Transferrin-Selenium (Gibco, 41400045). Type II AECs were seeded in tissue culture plates coated with gelatin-based coating solution (Cell Biologics, 6950).

**Immunoblotting**

Type II AECs were isolated from young and aged mice as described above. Cells were lysed using cold RIPA buffer (Sigma, 89900) containing 1% protease inhibitor cocktail (Sigma, P8340) and phosphatase inhibitor cocktail (Sigma, P5726). The lysate was then collected and centrifuged at 12,000 g for 15 minutes at 4°C. The supernatant was collected into a clean microcentrifuge tube and the pellet was discarded. Samples were denatured and reduced using NuPAGE LDS Sample Buffer (ThermoFisher, NP0007) and reducing agent (ThermoFisher, NP0009) according to manufacturer's instructions. Lysates were electrophoresed on 4-12% gradient Bis-Tris gel (ThermoFisher, NP0322BOX) and transferred to 0.2 micron PVDF membranes (ThermoFisher, IB401001). Blots were blocked for 1 hour at room temperature with PBS containing 0.01% Tween-20 and 5% BSA. Membranes were then incubated overnight at 4°C on an orbital shaker with the primary antibody. The membranes were blotted for p21 (Santa Cruz, sc-6246, 1:200) and the loading control GADPH (Cell Signaling, 14C10, 1:1000). After washing in PBS containing 0.01% Tween-20, membranes were incubated in an appropriate secondary antibody for 1 hour at room temperature (anti-mouse IgG, Abcam, ab205719, 1:10000; anti-Rabbit IgG, Abcam, ab205718, 1:10000). Blots were then wash with PBS containing 0.01% Tween-20, incubated with a chemiluminescent horseradish peroxidase substrate (Fisher, PI34577) for 5 minutes, and imaged using a BioRad ChemiDoc XRS+.

**Seahorse Assay**

Cells were seeded at a density of 2.5 x $10^4$ cells/well in a Seahorse XFe96 Cell Culture Microplate (Agilent) with completed RPMI 1640 media (see MH-S cell culture in Supplemental Experimental Procedures) containing 1µM $PGE_2$ or vehicle control. Cells were left to culture for 18 hours. Cells were then washed twice with Seahorse XF RPMI media (Agilent) supplemented with 1mM pyruvate, 2mM L-glutamine, and 10mM D-glucose; a final volume of 180µl was placed in each well. Cells were then incubated at 0% $CO_2$ and 37°C for 1 hour before being placed into a calibrated Seahorse XFe96 Analyzer (Agilent). Four basal readings were taken prior to the addition of the electron transport chain inhibitors in Agilent's mitostress test: 1.5µM oligomycin, 1µM FCCP, and 0.5µM rotenone and antimycin A (Agilent, cat# 103708). OCR and ECAR data were normalized to total cellular protein levels as measured by BCA assay (ThermoFisher, 23225). Seahorse experiments were repeated three times.

**BrdU in vivo**

BrdU (Biogems, 5911439) was given to mice in their drinking water *ad libitum* for a total of 7 days. BrdU was dissolved in the drinking water at a concentration of 0.8mg/ml and the BrdU water was refreshed every 48 hours.

**Transcriptomics Analysis**

Publicly available RNA-seq (GEO GSE134397) dataset of young and aged AM transcripts was downloaded from the Gene Expression Omnibus (GEO). Raw counts were processed in R (version 4.1.0) using edgeR (version 3.34.1) to generate TMM normalized counts (Robinson et al., 2010). The quasi-likelihood genewise dispersions were calculated using the glmQLFit function. Differentially expressed proteins between conditions were calculated by a quasi-likelihood negative binomial generalized log-linear model by using the glmQLFit function. False detection rate (FDR) *q* values (adjusted p-values) were used to correct for multiple comparisons. A value of FDR= 0.05 was used as a threshold for statistical significance. Publicly available scRNA-seq dataset of type I and type II AEC transcripts (GEO GSE113049) was downloaded from GEO and analyzed as described in Reimondy et al, 2019.

### Supplemental Figures

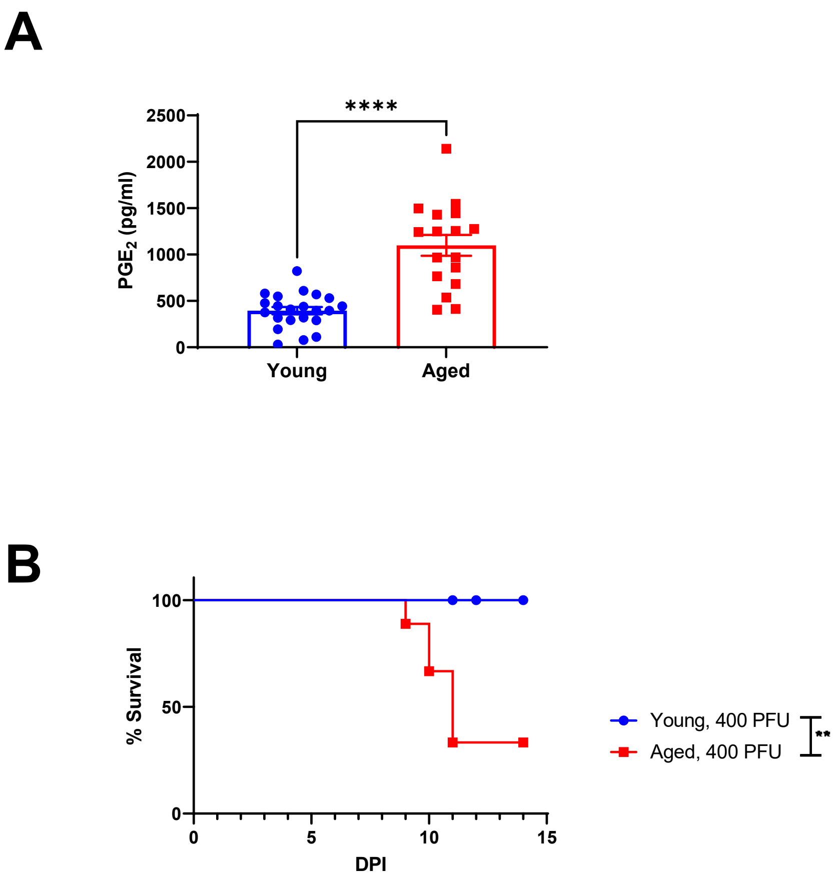


**Supplemental Figure 1:** (**A**) $PGE_2$ measured from the plasma of non-infected young (2-4M) and aged (18-22M) C57BL/6 female mice. Data analyzed by Mann-Whitney. Error bars represent SEM. Each point represents a biological replicate. (**B**) Percent survival of young (2-4 months) and aged (18-22 months) female C57BL/6 mice infected with 400 pfu of IAV i.n. n= 9 / group. Survival differences were statistically determined the Gehan-Breslow-Wilcoxon test**.** **p < 0.005, **** p< 0.0001

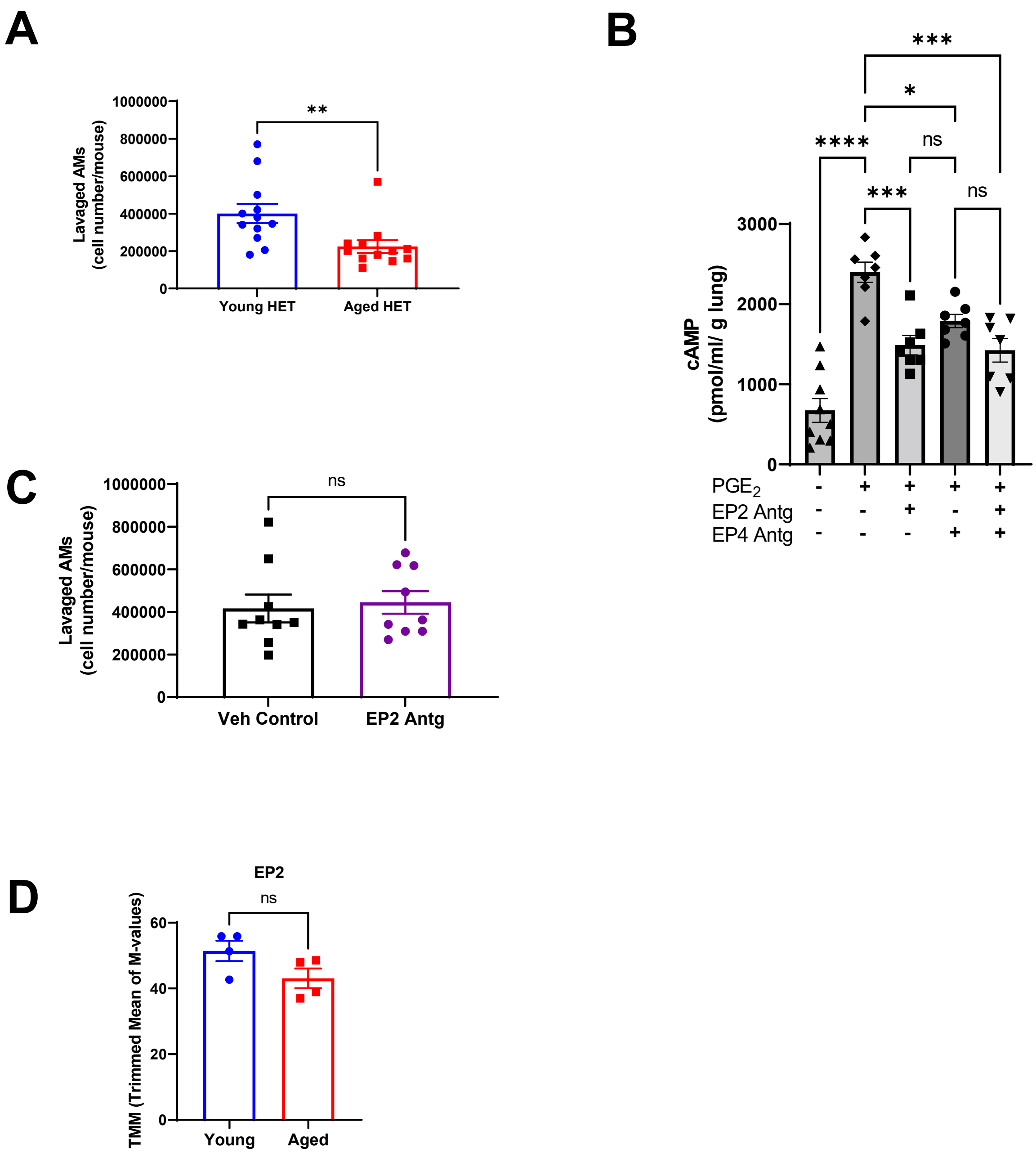


**Supplemental Figure 2**: (**A**) AMs in the BALF of young and aged male UM-HET3 mice were counted by flow cytometry. Statistical significance analyzed by Mann-Whitney. (**B**) Young (i.e., 2-4 months) C57BL/6 mice were given an i.p. injection of the EP2 antagonist and/or the EP4 antagonist or vehicle control followed by i.n. dosing of 15µg of $PGE_2$. Whole lungs were harvest 1 hour later. cAMP was measured in lung homogenate via ELISA. Statistical significance analyzed by ANOVA with Tukey post-hoc test. (**C**) Young (i.e., 2-4 months) C57BL/6 mice were given 7 daily i.p. injections of 10mg/kg EP2 antagonist. AMs (i.e., $CD45^+$ $CD11c^+$ $SiglecF^+$) were then collected through BAL and analyzed by flow cytometry. Total cell count of lavaged AMs were quantified. Statistical significance analyzed by Mann-Whitney. (**D**) Trimmed mean of M-values (TMM) normalized counts of EP2 of AMs sorted from young and aged C57BL/6 mice (GEO GSE134397). Statistical significance analyzed by Mann-Whitney. Error bars represent SEM. Each point represents a biological replicate. * $p < 0.05$, ** $p<0.005$, *** $p<0.0005$, **** $p<0.0001$

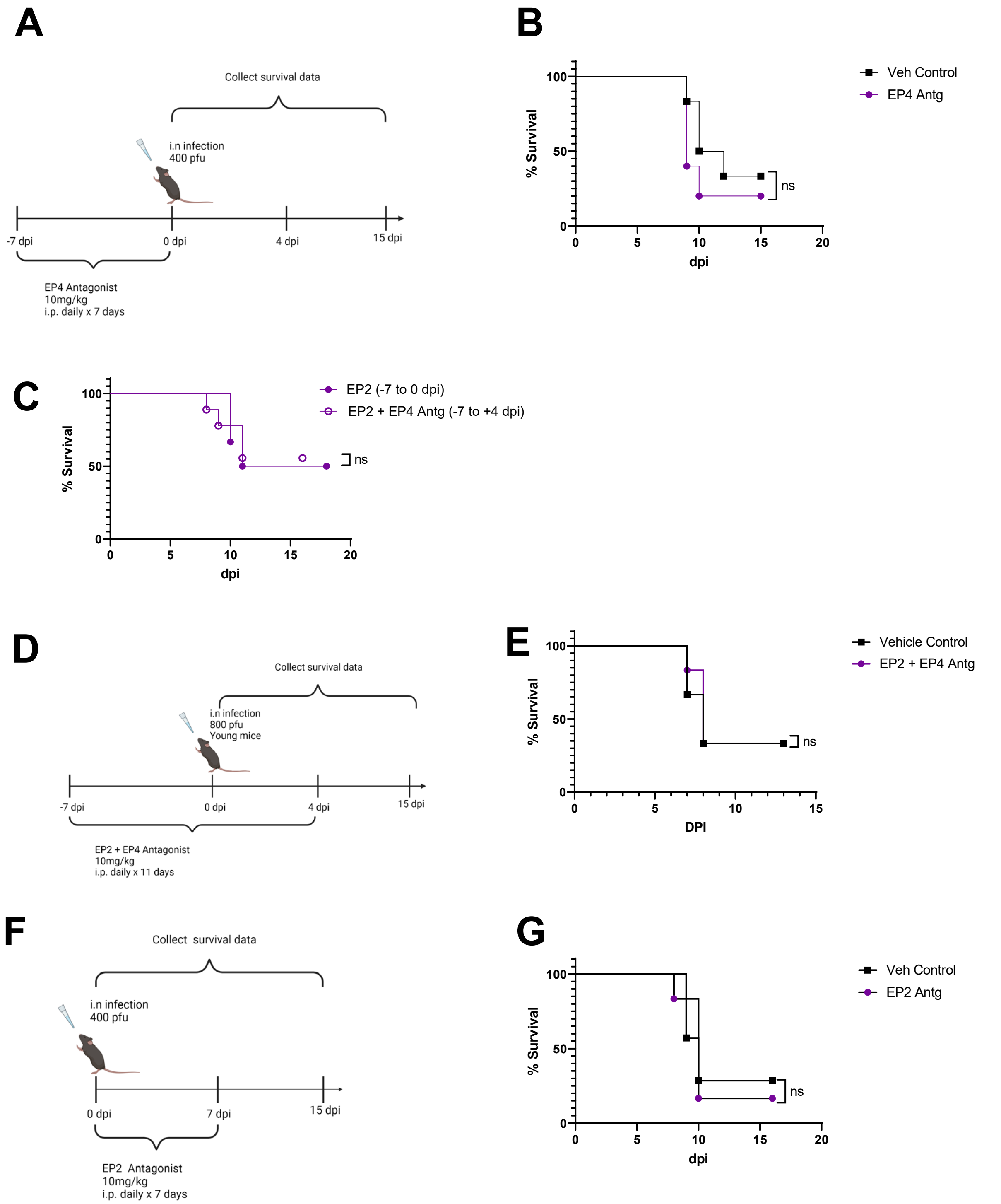


**Supplemental Figure 3:** (**A-B**) Aged C57BL/6 mice were prophylactically treated with the EP4 antagonist or vehicle control by daily i.p. injections for 1 week pre-infection as showin in (**A**). Surivival (**B**) were tracked daily. n = 5-7/group. (**C**) Percent survival of aged mice given the EP2 antagonist treatment as shown in Figure 3A and of aged mice given the EP2 and EP4 antagonist treatment as shown in Figure 3C. Survival data replicated from Figure 3B and Figure 3D. n = 6-9/group (**D-E**) Young female C57BL/6 mice were given the EP2 antagonist and EP4 antagonist or vehicle and infected with 800pfu of PR8 H1N1 as shown in (**D**). Survival of mice (**E**) were recorded daily. n = 6/ group. (**F-G**) Aged C57BL/6 mice were treated with the EP2 antagonist or vehicle control by daily i.p. injections for 1 week starting on the day of infection as shown in (**F**). Survival of mice (**G**) were tracked daily. n = 4-5 / group. Survival differences were statistically determined the Gehan-Breslow-Wilcoxon test**.** Schematics created in BioRender.

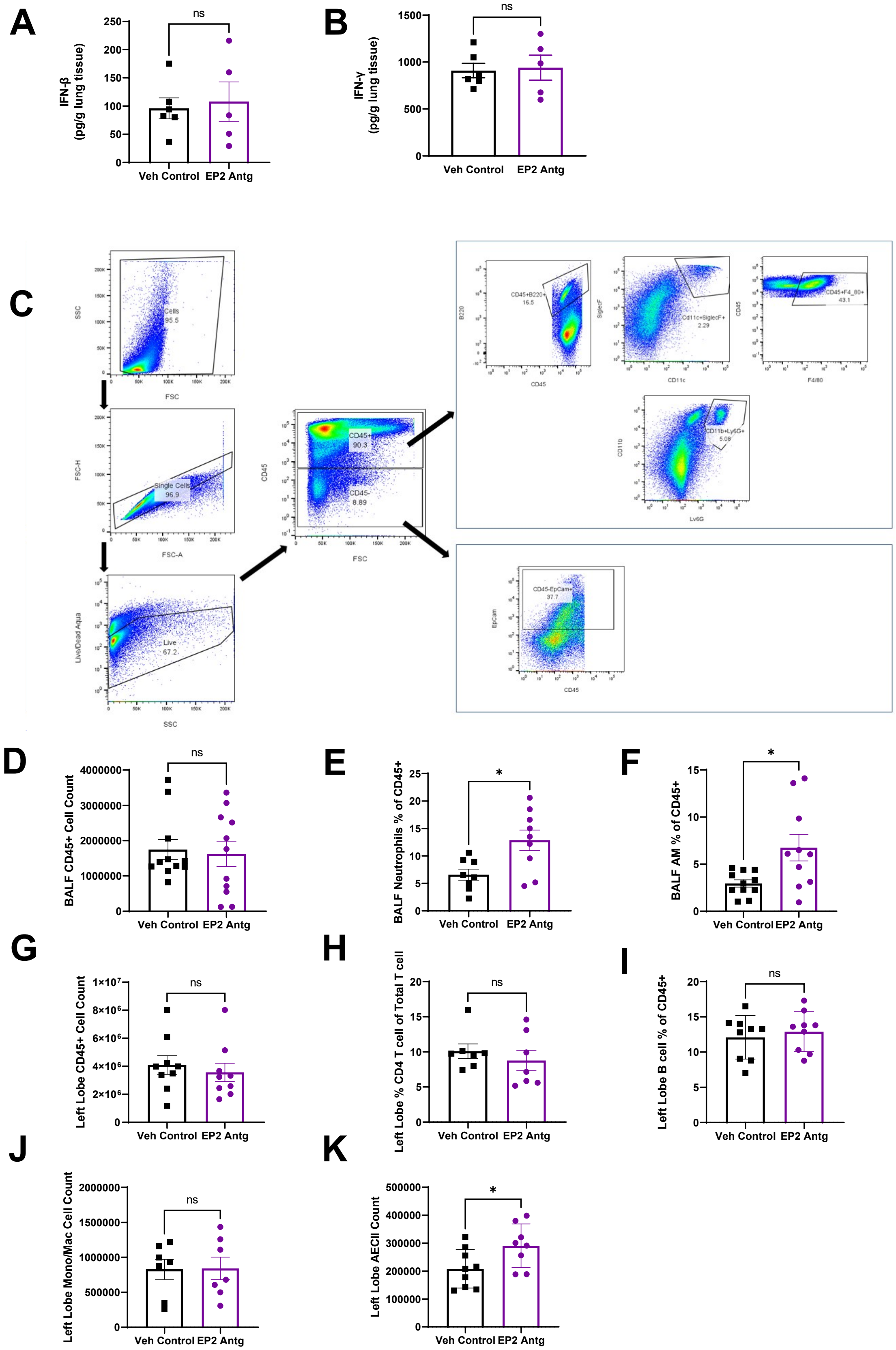
A
IFN-β (pg/g lung tissue)
ns
Veh Control
EP2 Antg
B
IFN-γ (pg/g lung tissue)
ns
Veh Control
EP2 Antg
C
Cells 95.5
Single Cells 96.9
Live 67.2
CD45+ 90.3
CD45- 8.89
CD45+B220+ 16.5
Cd11c+SiglecF+ 2.29
CD45+F4_80+ 43.1
CD11b+Ly6G+ 5.08
CD45-EpCam+ 37.7
D
BALF CD45+ Cell Count
ns
E
BALF Neutrophils % of CD45+
*
F
BALF AM % of CD45+
*
G
Left Lobe CD45+ Cell Count
ns
H
Left Lobe % CD4 T cell of Total T cell
ns
I
Left Lobe B cell % of CD45+
ns
J
Left Lobe Mono/Mac Cell Count
ns
K
Left Lobe AECII Count
*

**Supplemental Figure 4**: (**A-B**) IFN-β (**A**) and IFN-γ (**B**) measured from lung homogenate collected at 4 dpi of infected aged mice treated prophylactically with the EP2 antagonist or vehicle. Statistical significance analyzed by Mann-Whitney. (**C**) Gating strategy (**D-F**) Flow cytometric analysis of immune cells within the BALF at 9 dpi of vehicle or EP2 antagonist prophylactically treated aged C57BL/6 mice**.** (**D**) Total cell count of hematopoietic ($CD45^+$) (**E**) Percentage of neutrophils (i.e., $CD45^+$ $CD11b^+$ $Ly6G^+$) of total $CD45^+$ cells (**F**) Percentage of alveolar macrophages (i.e., $CD45^+$ $CD11c^+$ $SiglecF^+$) of total $CD45^+$ cells. (**G-I**) Flow cytometric analysis of immune cells within the left lung lobe at 9 dpi of vehicle or EP2 antagonist prophylactically treated aged C57BL/6 mice. (**G**) Total cell count of hematopoietic ($CD45^+$). (**H**) Percentage of $CD4^+$ T cells ($CD3^+$ $CD4^+$ $CD8^-$) of total T cells ($CD3^+$). (**I**) Percentage of B cells (i.e., $CD45^+$ $B220^+$) of total $CD45^+$ cells. (**J**) Total cell count of monocytes/macrophages (i.e., $CD45^+F4/80^+$). (**K)** Total type II AEC cell count (i.e.,$CD45^-$ $EpCAM^+$). Statistical significance analyzed by Mann-Whitney. Error bars represent SEM. Each point represents a biological replicate. *p<0.05.

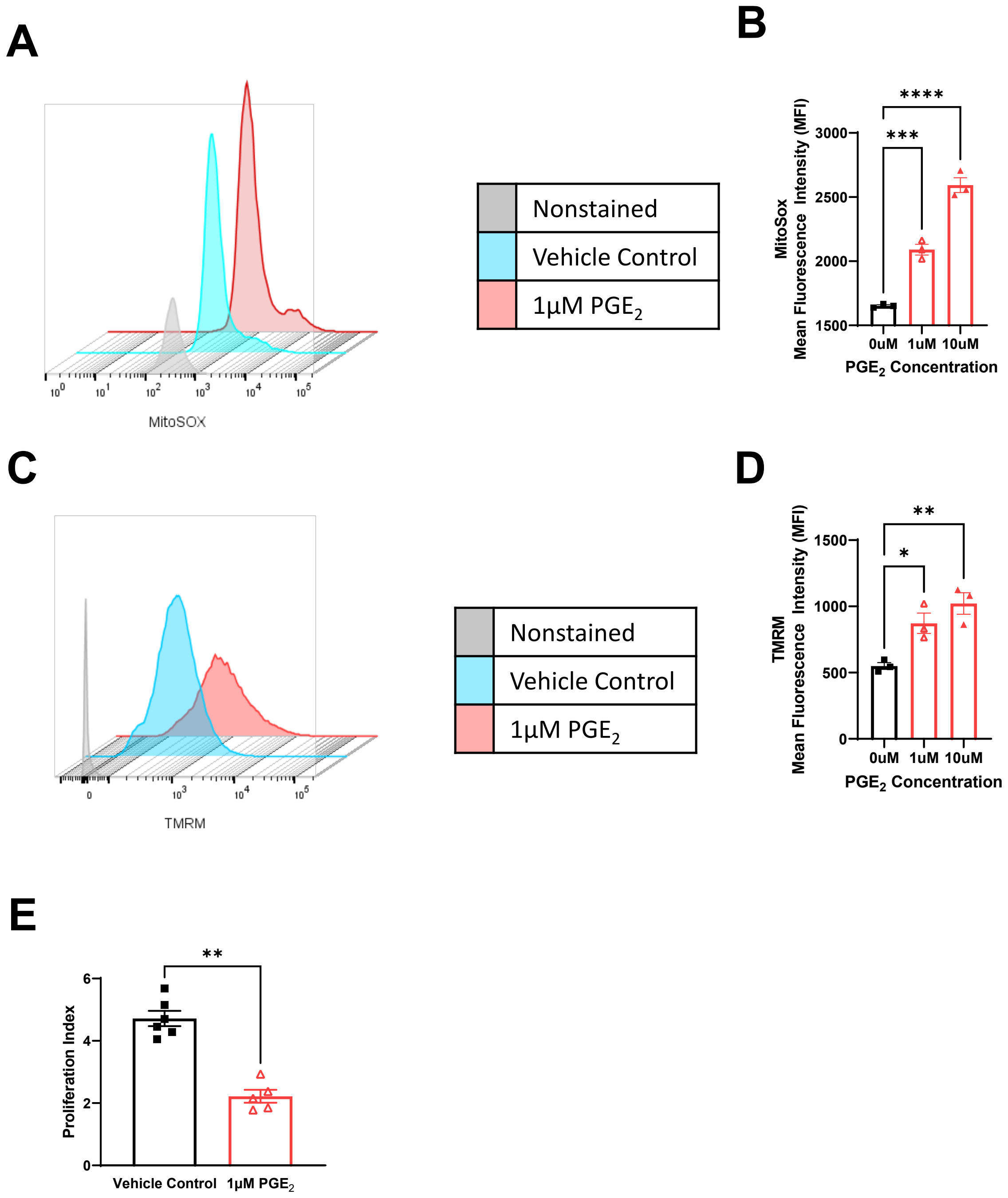


**Supplemental Figure 5**: MHS cells were cultured with varying concentrations of $PGE_2$ for 24 hours. Cells were than stained with (**A-B**) MitoSox and (**C-D**) TMRM and analyzed by flow cytometry. Mean fluorescence intensity was quantified. Statistical significance analyzed by ANOVA with Tukey post-hoc test. (**E**) Proliferation index of MH-S cells were obtained by counting cells under a hemocytometer and calculating proliferation index as (total cells) / (number of cells seeded). Statistical significance analyzed by Mann-Whitney. Error bars represent SEM. Each point represents a replicate. * $p < 0.05$, ** $p < 0.005$.

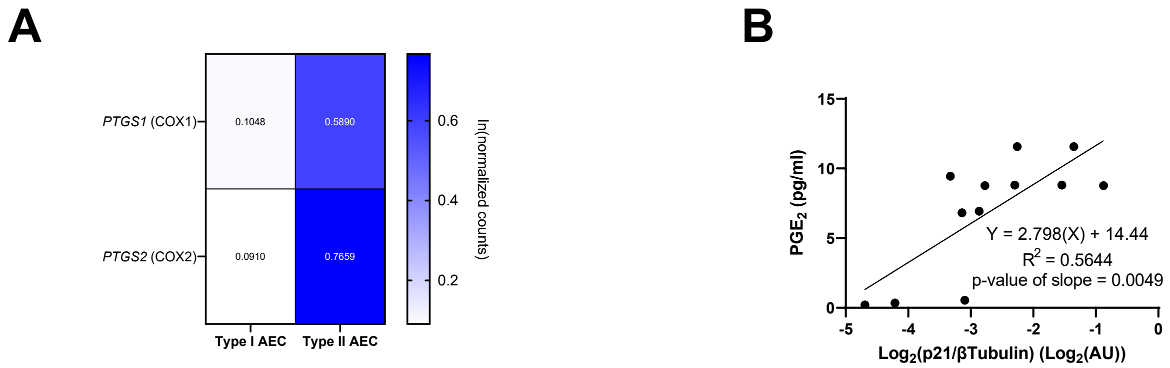


**Supplemental Figure 6**: (**A**) Heatmap of the ln(normalized counts) of *PTGS1* and *PTGS2* (gene products are COX1 and COX2, respectively) of type I and type II AECs (GEO GSE113049). (**B**) p21 expression level correlates with $PGE_2$ production in type II AECs. Type II AECs from young mice were isolated and irradiated with 0, 5, 10, and 15 Gy to induced senescence as measured by western blotting for p21. $PGE_2$ was measured from the culture medium by ELISA. $PGE_2$ levels was plotted against log-transformed, β-tubulin normalized, p21 expression and a linear regression analysis was performed. The p-value of the slope was calculated based on the null hypothesis of slope = 0 and the alternative hypothesis of slope ≠ 0.
